# Structure induces computational function in networks with diverse types of spiking neurons

**DOI:** 10.1101/2021.05.18.444689

**Authors:** Christoph Stöckl, Dominik Lang, Wolfgang Maass

## Abstract

Nature endows networks of spiking neurons in the brain with innate computing capabilities. But it has remained an open problem how the genome achieves that. Experimental data imply that the genome encodes synaptic connection probabilities between neurons depending on their genetic types and spatial distance. We show that this low-dimensional parameterization suffices for programming fundamental computing capabilities into networks of spiking neurons. However, this method is only effective if the network employs a substantial number of different neuron types. This provides an intriguing answer to the open question why the brain employs so many neuron types, many more than were used so far in neural network models. Neural networks whose computational function is induced through their connectivity structure, rather than through synaptic plasticity, are distinguished by short wire length and robustness to weight perturbations. These neural networks features are not only essential for the brain, but also for energy-efficient neuromorphic hardware.

**Significance statement:** Fundamental computing capabilities of neural networks in the brain are innate, i.e., they do not depend on experience-dependent plasticity. Examples are the capability to recognize odors of poisonous food, and the capability to stand up and walk right after birth. But it has remained unknown how the genetic code can achieve that. A prominent aspect of neural networks of the brain that is under genetic control is the connection probability between neurons of different types. We show that this low-dimensional code suffices for inducing substantial innate computing capabilities in neural networks, provided they have –like the brain– a fair number of different neuron types. Hence under this condition structure can induce computational function in neural networks.

## 1 Introduction

The common paradigm for bringing computational function into neural networks, including models for neural networks of the brain, is to tune their very large number of synaptic weights by a learning process, starting from a tabula rasa initial state. This typically requires very large numbers of training examples, which are for many tasks not readily available. Nature has invented a powerful alternative: The genetic code endows neural networks of the brain with an exquisite structure that induces numerous computational capabilities without a need for experience-dependent plasticity, see Zador, 2019 for a review, and (Apfelbach et al., 2005, Yilmaz and Meister, 2013, Tinbergen, 2020, Weber and Hoekstra, 2009, Metz et al., 2017, Langston et al., 2010, Mckone, Crookes, and Kanwisher, 2009) for experimental data. In fact, innate functional capabilities, such as avoidance of poisonous food and the capability to stand up and walk right after birth, are in many cases crucial for survival. However, it has remained an open problem how the genetic code achieves that. Nature must have found a way to encode the computational function through a low-dimensional parametrization, rather than by encoding individual synaptic weights, since even the human genome contains only about 1 GB of information (Zador, 2019). We show that known genetically encoded structural properties of cortical microcircuits provide a solution to this problem. Experimental data on cortical microcircuits, such as Markram et al., 2015 and Billeh et al., 2020, prove that the genetic code determines connection probabilities in terms of the genetic type of the pre- and postsynaptic neuron and their spatial distance. We show that these structural features of networks of spiking neurons suffice for inducing specific computational functions. This insight provides simultaneously an answer to another open question: Why the brain employs so many neuron types, substantially more than we have commonly considered in neural network models.

We base our answers to these open problems on a new type of generative model, a probabilistic skeleton. Neural networks that are generated by a probabilistic skeleton share a number of salient statistical features with neural networks in the brain that are under genetic control, such as the number and prevalence of neuron types, and connection probabilities in terms of these neuron types.

Probabilistic skeletons generate just the architectures of neural networks, hence these can in principle employ any kinds of computational units. We focus here on networks that consist of excitatory and inhibitory spiking neurons (RSNNs). These are of particular interest for modelling neural networks of the brain because the activity of these units can be related directly to neural recordings from the brain, especially if the RSNN operates in an event-driven sparse firing regime where the timing of spikes can be used to encode salient information. However, it has turned out to be difficult to endow RSNNs with powerful computational capabilities through training, in particular if one wants that they operate in a spare firing regime. Hence inducing function through structure is a particularly desirable tool for RSNNs. Producing computationally powerful RSNN models that operate in a sparse firing regime is also of interest in the quest to design more energy-efficient computing hardware for AI, because hardware implementations of sparsely active RSNNs tend to consume substantially less energy than customary digital computing hardware (Plank et al., 2022).

We will first define the concept of a probabilistic skeleton, and then show that they suffice to induce specific computing capabilities in RSNN. In particular, we consider examples for generic 2D computing capabilities of laminar cortical microcircuits, the capability to recognize particular spike patterns, and to carry out a generic motor control task. Finally, we will elucidate principles of this new method to generate network function through network structure.

## 2 Results

### 2.1 Probabilistic skeletons provide a mathematical model for aspects of network generation that are under genetic control

Current models of cortical microcircuits (Markram et al., 2015; Billeh et al., 2020) are based on two types of data: A set of neuron types -estimated to be well over 100 within a single neocortical area (Tasic et al., 2018) - and a table of connection probabilities for any pair of neuron types as in panel A of Fig. 4 in (Billeh et al., 2020), which is reproduced here as Fig. 1a. The entries of this table provide base connection probabilities that are valid if the somata have a horizontal distance of at most 75*μm.* If the horizontal distance is larger, these base connection probabilities are multiplied with an exponentially decaying function of their distance. Examples of such functions are shown in panel C of Fig. 4 in (Billeh et al., 2020), reproduced here as Fig. 1b.

**Figure 1:**
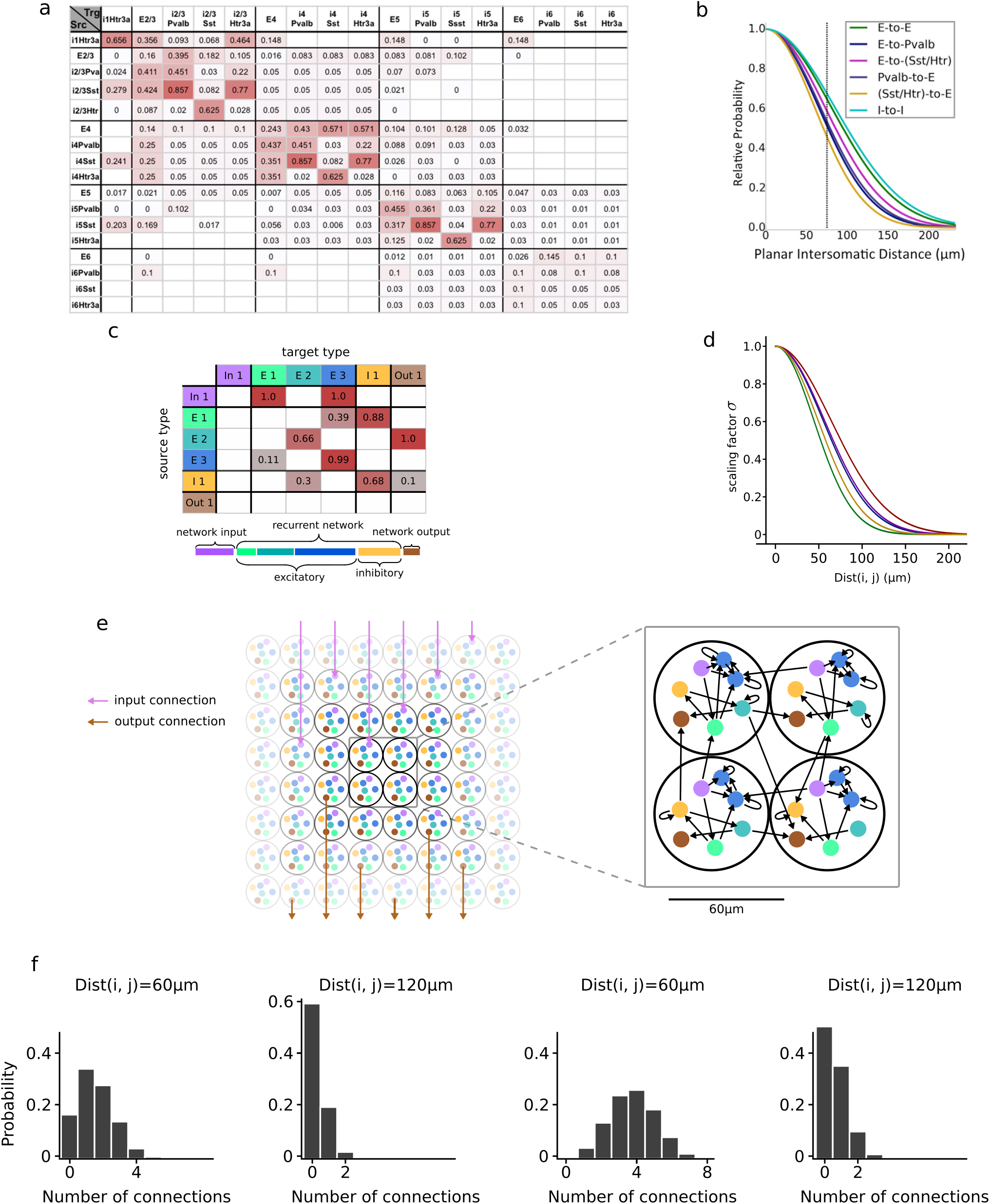
Illustration of a probabilistic skeleton and the process by which RSNNs are generated from it. **a** Base connection probabilities between 17 types of neurons in mouse V1 (reproduced from (Billeh et al., 2020)). White table cells indicate unknown values. **b** Scaling of connection probabilities with the horizontal distance of their somata for mouse V1 (reproduced from (Billeh et al., 2020)). **c** Top: Sample base connection probability table of a probabilistic skeleton for the case of *K* = 6 neuron types. White table cells indicate here that the corresponding base connection probability has value 0. Rows and columns labeled “in” refer to input neuron types, the label “out” refers to output neuron types, “E” (“I”) to the other excitatory (inhibitory) types of neurons. Bottom: Prevalence-bar of a probabilistic skeleton. Its length defines the number *M* of neurons in a minicolumn. **d** Examples of distance-dependent scaling functions, with slightly different values of *σ* in equation (4). These functions turned out to work well for the computing tasks that we considered. **e** Illustration of the uniform distributions of neurons of all types over a 2D sheet for the generation of an RSNN from a probabilistic skeleton. Each disc can be seen as 2D projection of a stereotypical minicolumn in the neocortical sheet. Sample arrows in purple indicate for some of them external inputs (that arrive at the purple neurons in each disc). Network outputs from brown neurons in all discs are indicated by brown arrows from a random sample. Synaptic connections are not restricted to neurons in the same or neighboring disc, the blow-up might suggest, but are drawn according to a distribution as shown in b. **f** Examples for binomial distributions from which the number *m_ij_* of synaptic connections (and hence the effective connection strength) from a neuron *i* of type *I* to a neuron *j* of type *J* are drawn for the case *p_I→J_* = 0.35 (two panels on the left) and *p_I→J_* = 0.85 (two panels on the right), each for two different values of the spatial distance Dist(*i, j*) between their somata.

A probabilistic skeleton is a rigorous generative model for this indirect style of encoding network architectures. It specifies the number K of neuron types, the prevalence of each neuron type (see the lower part of Fig. 1c), and base connection probabilities in dependence of neuron types (see the upper part of Fig. 1c). In addition, it specifies a parameter *σ* that scales the exponential decay of connection probabilities with the lateral distance between the somata according to equation 4 in Methods; see Fig. 1d for samples of exponential decays that were useful for tasks that we considered (although the precise shapes had little impact). A probabilistic skeleton does not specify the values of individual synaptic weights, but it specifies three parameters *w_in_, w_E_, w_I_* that define the weight of each synaptic connection from an input neuron (i.e., a neuron that also receives synaptic input from outside of the RSNN), from an excitatory neuron that is not an input neuron, and from an inhibitory neuron that is not an input neuron. Input and output neurons (projection neurons) are from separate neuron types, and are assumed to be embedded into the RSNN (Fig. 1g). Neurons in the neocortex that are synaptically connected are usually connected by multiple synaptic connections, see e.g. Fig. 7A of (Markram et al., 2015). Hence we draw for each pair *i, j* of neurons not just once, but *S* times from the corresponding connection probability, see equation 4. We used S = 8 in the experiments that are reported here, but the exact value had little impact. The multiplicity *m_ij_* of synaptic connections between two neurons induces some differentiation in the effective strength (weight) by which two neurons are connected: One multiplies the corresponding parameter *w_in_, w_E_, w_I_* that determines the uniform synaptic weight of all such synapses with the actual number of synaptic connections between the two neurons. Hence the effective strength of the connection between two neurons is drawn from binomial distributions for their connection probability, in dependence of their types and distance (Fig. 1f).

To sample a neural network from a probabilistic skeleton, one needs to specify its number of neurons *N*. One also needs to specify their spatial positions, because their distances are relevant for their connection probabilities. Actually, in the neocortex primarily the horizontal (lateral) distance within the 2D neocortical sheet is relevant for that. Therefore it suffices to distribute the neurons of each type uniformly over a 2D sheet. A convenient method for doing that is to let the 2D sheet consist of a grid of discs that each contain the same number of neurons with the specified prevalence of different neuron types, see Fig. 1e for an illustration (for tasks with small numbers of input or output neurons these were placed into selected subsets of the discs). For measuring horizontal distances between neurons we assume for simplicity that the neurons are always positioned at the center of a disc. In terms of neural anatomy each disc can be seen as a 2D projection of a minicolumn. It is well-known that the neocortical sheet is made up of stereotypical minicolumns of diameter around 60 μm that extend vertically across all neocortical layers, and each contains a representative sample of together 80 - 120 neurons of all types (Mountcastle, 1998, Cruz et al., 2005, DeFelipe, 2015).

Since a probabilistic skeleton only captures aspects of the architecture of neocortical neural networks that are under genetic control, one can use this concept to examine the impact of genetically encoded architectural features on computational properties of a neural network. If a probabilistic skeleton endows with high probability its neural network samples with a specific computing capability, this computing capability can be argued to be within the reach of genetic control (i.e., “innate”).

We used evolution strategies (Schaul, Glasmachers, and Schmidhuber, 2011) to optimize the parameters of a probabilistic skeleton for a given computational task, see Fig. 7 for an illustration. Note that a fitness function that measures the computational performance of RSNN samples from a probabilistic skeleton is not differentiable because RSNNs are sampled from it using a stochastic process.

### 2.2 Generic 2D computing capabilities of cortical microcircuits

The neocortex forms a 2D sheet composed of several parallel laminae or layers that each consist of different types of neurons (Mountcastle, 1998; Harris and Shepherd, 2015; Billeh et al., 2020). Sensory input streams and outputs from other brain areas also have a clear 2D structure, and they are mapped topographically onto specific layers of generic laminar cortical microcircuits. Of special importance is the capability to compare 2D patterns that arrive from lower brain areas and higher brain areas in hierarchical brain networks (Vezoli et al., 2021), typically with some time-gap in between. Hence we are focusing here on the task to decide whether two 2D patterns that arrive sequentially, with varying delays between them, in different 2D input layers, are similar or not (see Fig. 2a). We demanded that a population of output neurons fires in the case of a non-match, in analogy to the error-reporting neurons found in the neocortex (Keller and Mrsic-Flogel, 2018). We found that a probabilistic skeleton with 7 recurrent neuron types, i.e., neuron types that are not marked as input or output neurons can solve this task convincingly (see Fig. 2). The resulting network connectivity of an RSNN sample is shown in section 1.1 in the Suppl. (Fig. S1), plotted in the same style (as chord diagram) as experimental data on network connectivity of the neocortex in Fig. 7C of Markram et al., 2015. The RSNN sample achieved an accuracy of 91.5%. Two trials, one of which should be judged as a “match” on the left, and a non-match trial on the right, are shown in Fig. 2c. Further trial input patterns can be seen in Fig. 2d and e.

**Figure 2:**
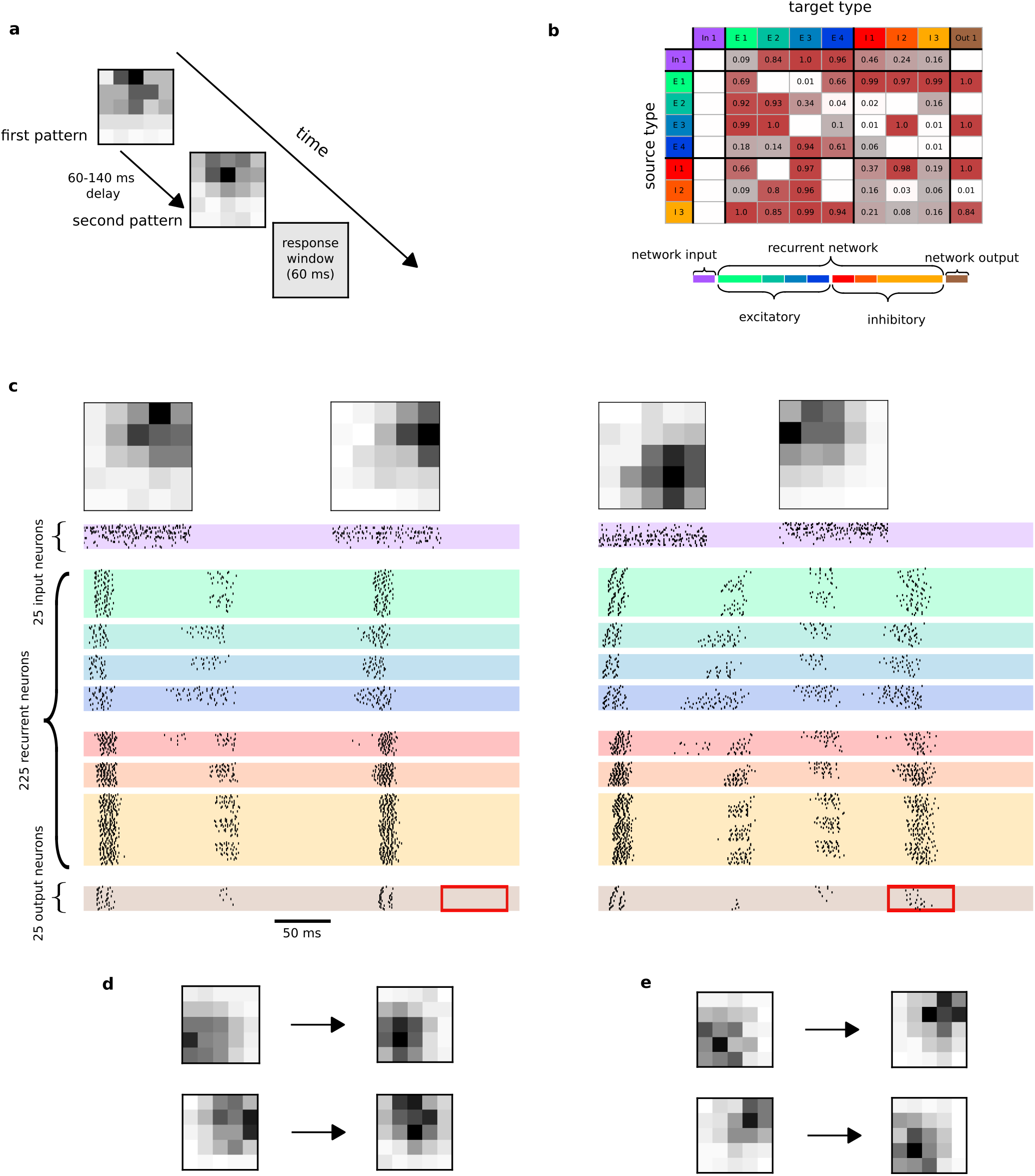
Induction of a fundamental computational capability of laminar cortical microcircuits, to decide whether the sequentially presented 2D network inputs are similar, through network structure, i.e., through a probabilistic skeleton. **a** Temporal structure of the task. **b** A probabilistic skeleton that induces high RSNN performance on this task. **c** Network inputs and firing activity of an RSNN sample from this probabilistic skeleton for two trials, with varying delay between the two input patterns. In the first trial the two patterns are correctly judged by the RSNN to be similar, indicated by withholding of firing of output neurons (shown at the bottom) during the response window (indicated by red frame). In the second trial the two input patterns were correctly judged by the network to be dissimilar. Note that information about the first pattern had to be retained within the network until the second pattern arrived. This working memory aspect was nontrivial because each pattern consisted of 25 gray values. Persistent firing of neurons of type E1 emerged as a mechanism for that. **d** Two further correctly classified samples of matching input patterns of activity. The delay between the first pair was 62ms while the delay between the second pair was 122ms. **e** Two additional correctly classified samples for non-matching input patterns. The delay between the first pair was 64ms and the delay between the second pair was 133 ms.

A related task that is also arguably central to innate computing capabilities of cortical microcircuits is the identification of coincidences in two 2D input streams, that could arrive from different sensory areas, or from a higher and a lower cortical area -one indicating spatial attention and another visual input. An essential sub-computation, that is arguably innate, is to mark those locations where both 2D input patterns have substantial activity. This computational capability can also be induced by a probabilistic skeleton, using just 121 parameters, see Fig. S2 e - g.

Another innate computing capability is likely to be contrast enhancement, which can also be induced by a probabilistic skeleton (see section 1.2 in the Suppl. and Fig. S2 a - b and Fig. S3).

Altogether we found that fundamental 2D computing operations that are arguably central for computational operations in generic cortical microcircuits can be induced through genetically encoded network structure.

### 2.3 Innate recognition of particular stimuli

We know that numerous species have innate capabilities to recognize particular stimuli, such as odors and/or views of poisonous food and predators. Since such stimuli arrive in the brain in the form of specific spatio-temporal spike patterns, we need to understand how the genetic code can install in RSNNs the capability to recognize specific spatiotemporal patterns, such as those depicted in Fig. 3a. We show that templates for any such patterns can be encoded in features of RSNN structures that are under genetic control. We fixed two randomly generated spatio-temporal spike pattern templates, and generated by adding, deleting, and shifting spikes noisy variations of these templates as inputs of class 1 and 2. We also created a class 3 of distractor patterns that were not similar to any of the two frozen template patterns but used the same firing rates as these. Three output neuron types were selected that were supposed the class membership of patterns from any of these 3 classes of spatio-temporal spike patterns. A probabilistic skeleton with 9 types of neurons (besides input- and output neuron types) with altogether 157 parameters (see Fig. 3b) is capable of achieving 91% accuracy on this task. A sample run of an RSNN sample from this skeleton, consisting of 144 neurons, for a pattern of class 1 is shown in Fig. 3c.

**Figure 3:**
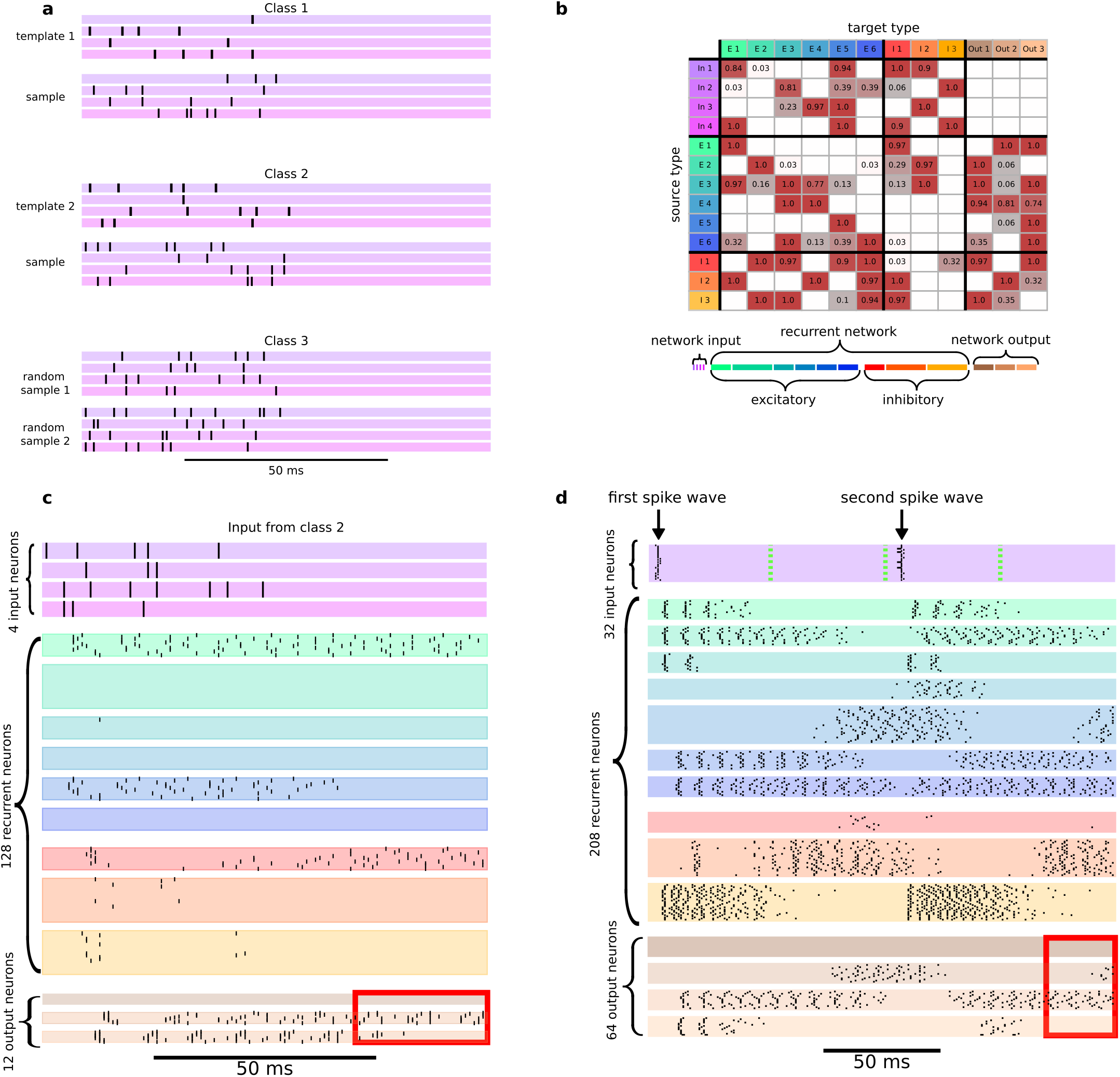
Innate spike pattern classification capability. **a** Two samples from each of the three classes of spike input patterns. The first two classes consist of variations of specific but arbitrarily chosen spike patterns, the third class consists of distractor spike patterns with the same firing rates. **b** Optimized probabilistic skeleton for this task. **c** Firing activity is shown for all neurons of RSNN samples with 144 neurons, in sample trials for spike inputs from class 2. The 30 ms time window during which the network decision is expected is indicated by the red frame at the bottom of the spike rasters. **d** Firing activity on the temporal pattern classification task, where the model tries to classify in which of the four time bins (50ms each) the second spike wave has arrived.

**Figure 4:**
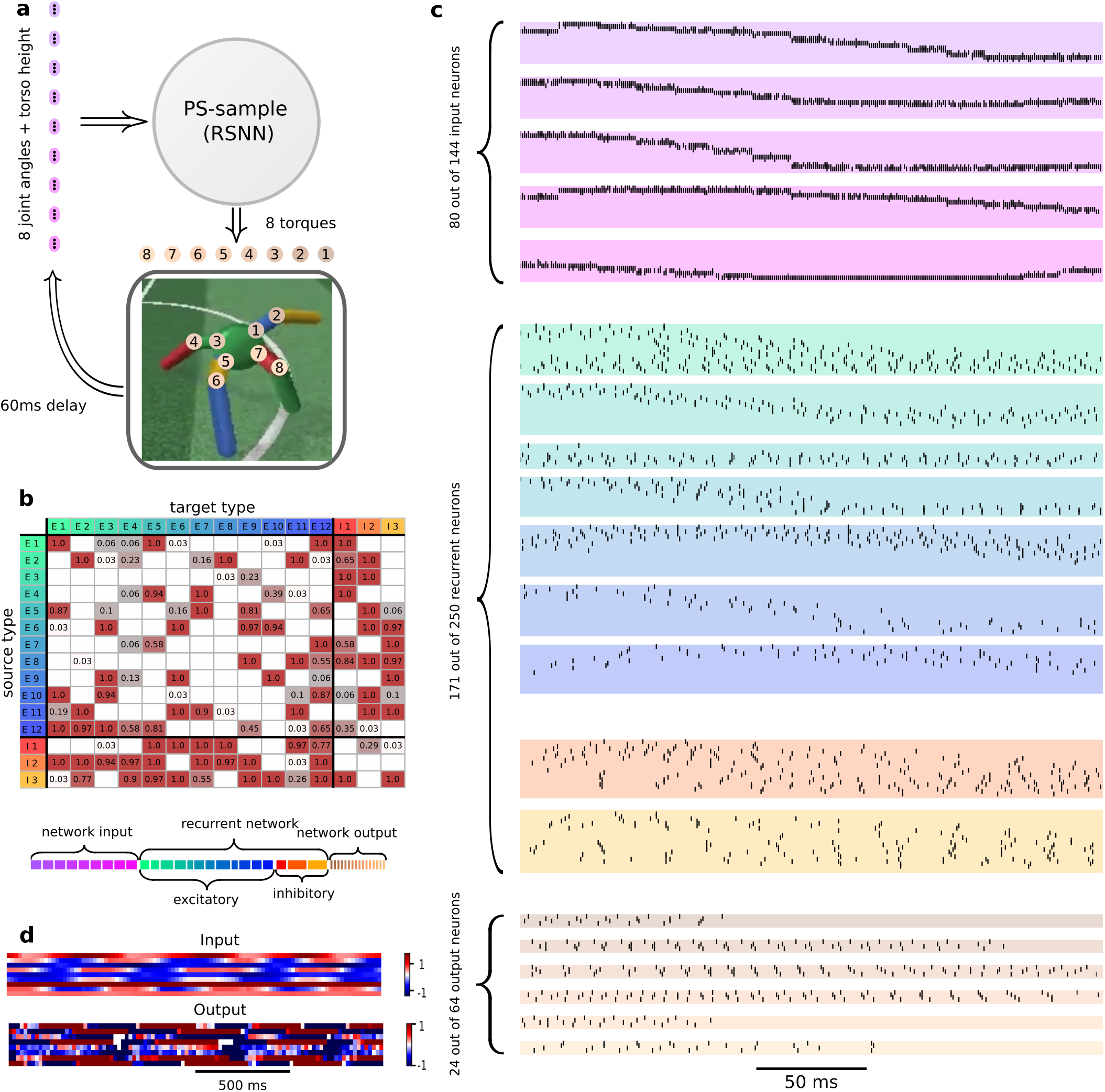
Example for innate motor control capability through a probabilistic skeleton. **a** System architecture, indicating network inputs and outputs, as well as the 8 joints that are controlled by the RSNN outputs. **b** Probabilistic skeleton for solving this motor control task (base connection probabilities for its numerous input- and output neuron types are shown in the Suppl., Fig S9) **c** Spike raster of an RSNN sample with 458 neurons drawn from this probabilistic skeleton. Population coding of the 9 continuousvalued input variables induced spatially structured firing activity in most of the neuron types. **d** Sample dynamics of input and output variables of the RSNN controller on a larger time scale.

One may wonder whether also the capability to distinguish purely temporal patterns can be induced through the structure of RSNNs. We used as test inputs two waves of input spikes with temporal distances from 1 to 200ms, where the task is to classify in which of the four time bins of 50ms each the second spike wave arrived. A probabilistic skeleton with just 10 neuron types (besides input and output neuron types) is capable of achieving an accuracy of 97% on this task. A typical spike raster of an RSNN sample is shown in Fig. 3d. One sees that temporal distances between the two waves of input spikes and the decision time are bridged by persistent activity in specific recurrent neuron types. Further spike rasters can be found in Fig. S6, S7 and S8. Altogether the results of this section demonstrate that also all-important capability of RSNNs to recognize and compute with temporal differences between spikes can be engraved into them through their structure.

The time scale of this task is in the range of behavioral responses, but a clever network organization is needed in order to enable a network of standard neuron and synapse models to discern and classify such fairly large time differences up to 200ms, and to produce the decision at a specific time after the onset of a trial, without an external clock or prompt.

### 2.4 A probabilistic skeleton can endow neural networks with innate motor control capability

Innate rudimentary motor control capability, for example, to stand up and walk right after birth, is essential for survival in many species. In contrast to the previously discussed computing tasks, biological motor control requires a transformation of multiple spike-input streams -that represent sensory inputs and feedback-into multiple time-varying spike output streams that control muscles in a closed loop, hence in real-time. We chose a standard benchmark task for motor control: Enabling a quadruped (“ant”) to walk by controlling the 8 joints of its 4 legs through a suitable stream of torques. The RSNN controller received 9 spike input streams that encoded -with a delay of 60 ms to make the task more challenging and biologically realistic-through population coding 9 dynamically varying variables: The angles of the 8 joints as well as the vertical position of the torso, see Fig. 4a. Further information about population coding can be found in the Suppl. in section 1.9 and in Fig. S5. We found that a probabilistic skeleton with just 15 types of neurons in the recurrent network, specified by 635 parameters, see Fig. 4b, is able to encode this motor control capability. We refer to movie of the ant locomotion^1^ for the resulting locomotion of the quadruped when its joints were controlled by an RSNN sample from this probabilistic skeleton. One can see in the input/output sample shown in Fig. 4d that the computational transformation which this task requires is quite complex. A sample spike raster of this RSNN in Fig. 4c shows that the population coding of the continuous-valued input variables induced a rather complex spatial dynamics of firing activity in most of the neuron types.

We employed RSNN samples from the probabilistic skeleton whose recurrent network consisted of 250 neurons. Direct tuning of their synaptic weights for this control task would result in a 114, 500 dimensional encoding of the control algorithm. The compressed encoding of the control strategy into just 635 parameters enhanced, as expected, the robustness of the RSNN controller: After randomly deleting 30% of the recurrent and output neurons of the RSNN, it was still able to control the ant locomotion, although the ant was walking somewhat slower, see (Movie of ant after 30% deletion)^2^. Altogether we have seen in this section that also demanding real-time computing capabilities in a closed loop with the environment, as required for locomotion, can be encoded in a relatively low-dimensional parameter space and induced in RSNNs through their structure.

### 2.5 Principles of structure-induced network function

#### Principle 1: A qualitative jump in computational performance of RSNN-samples occurs for many tasks when substantially more than 2 types of neurons are allowed

Fig. 5a for the tasks considered in sections 2.2 and 2.3. Also the probabilistic skeleton that controls locomotion of a quadruped (section 2.4) requires substantially more than 2 neuron types.

**Figure 5:**
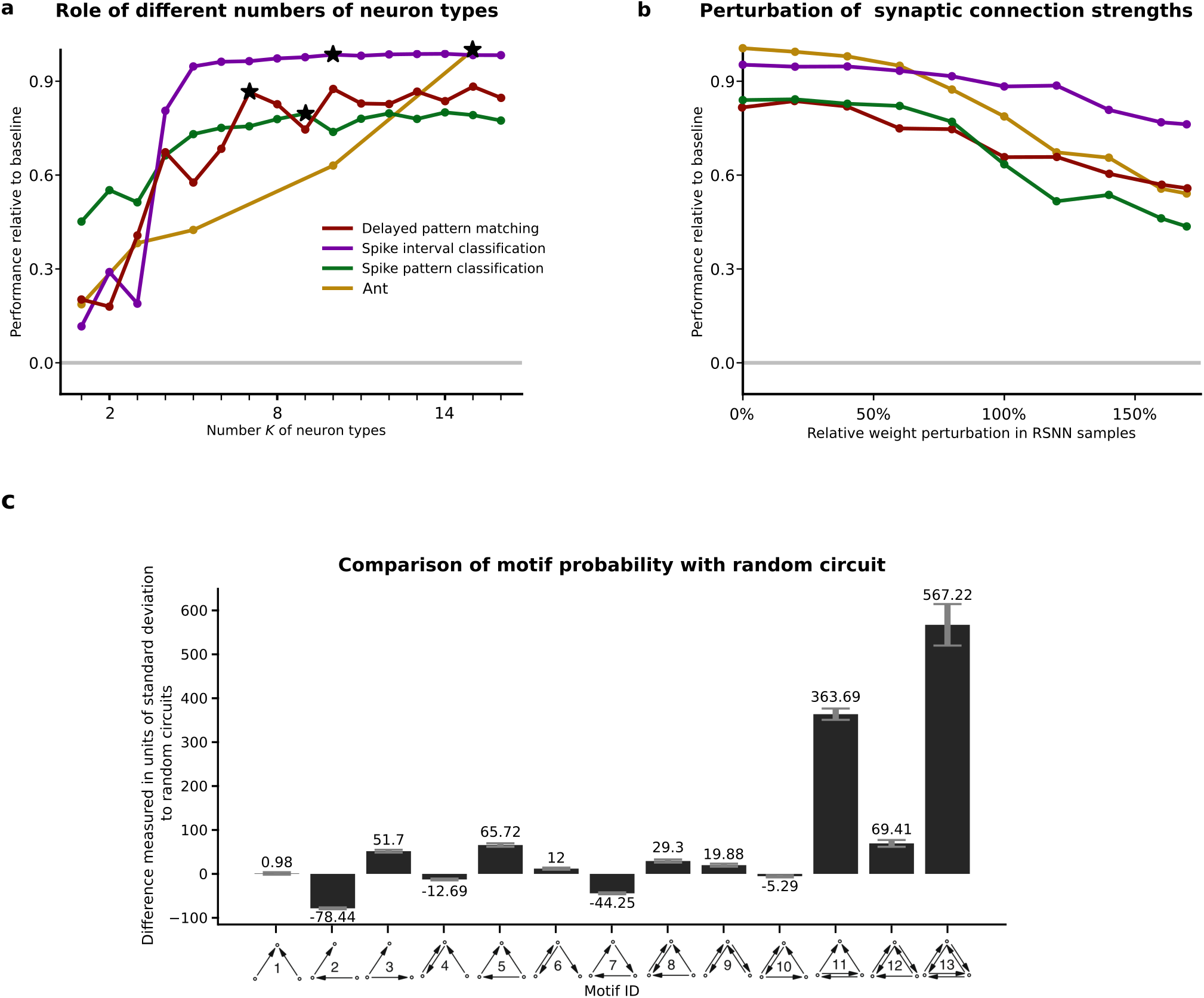
Empirical results underpinning Principles 1.- 3. **a** Performance achieved by optimizing probabilistic skeletons with different numbers of recurrent neuron types. One sees a qualitative jump in computational performance when substantially more than 2 neuron types are considered. Black stars mark the numbers of neuron types of the probabilistic skeletons that were discussed for each task in preceding sections. Performance of RSNN was measured relative to random guessing as a common baseline for the tasks considered in sections 2.2 and 2.3, see Methods for details. **b** The computational performance of the RSNNs degrades gracefully when the strengths of synaptic connections, defined by the number of synaptic connections between two neurons, is randomly perturbed. **c** Comparison of the frequency of three neuron network motifs occurrence in a generic RSNN sampled from a probabilistic skeleton (the one from Fig. 4) and in a randomly connected graph with the same number of nodes and directed edges.

#### Principle 2: Structure-induced network function is inherently robust to noise in synaptic strength

Different strengths of synaptic connections arise in a RSNN generated by a probabilistic skeleton from multiple synaptic connections between two neurons. One sees that the network performance is quite robust to random perturbations of these synaptic strengths, as can be seen in Fig. 5b. This may explain why the performance of the RSNNs in the neocortex is little affected by continuously ongoing spine motility (Yasumatsu et al., 2008; Holtmaat and Svoboda, 2009)

#### Principle 3: Probabilistic skeletons with an exponential decay of connection probabilities generate RSNNs whose number of synapses and total wire length grow just linearly with the number of neurons, and which have more strongly interconnected clusters of neurons than randomly connected graphs

The over-expression of strongly interconnected network motifs in RSNNs that are sampled from a probabilistic skeleton arises from the fact that the majority of synaptically connected neurons in such an RSNN have small distance. This strongly increases the chance of having also a synaptic connection in the opposite direction, and also favors the emergence of stongly interconnected clusters of neurons (see Fig. 5 c for a sample). This is consistent with experimental data on the connectivity structure of neural networks in the cortex, where one finds a similar over-expression of strongly interconnected groups of neurons Song et al., 2005 and Perin, Berger, and Markram, 2011.

We refer to Suppl. sections 1.7 and 1.8 for concrete estimates of the expected number of synapses and wire length per neuron in RSNNs that are sampled from a probabilistic skeleton.

#### Principle 4: Activity patterns and computations in network samples from probabilistic skeletons can be arbitrarily complex

Any cellular automaton, in fact more powerful versions where cells can have connections not only to the immediately adjacent cells but to a larger neighborhood of cells, arise as special cases of networks that can be generated from a probabilistic skeleton. This is demonstrated in Fig. 6 for a particularly well-known cellular automaton, the Game-of-Life. It has attracted substantial interest because it can emulate any Turing machine, and hence any digital algorithm (Soare, 2016). Each cell of this cellular automaton can assume two states: Dead or alive. It is alive at some time step if and only if either exactly 3 of its 8 neighbors in a 2D grid (one counts here also neighbors that just share a corner) were alive at the preceding time step, or if the cell itself and exactly 2 of its neighbors were alive at the preceding time step. In the RSNN that arises from the probabilistic skeleton with 5 neuron types shown in Fig. 6a, a neuron of type E1 indicates at every second time step through firing or not firing whether the cell of the cellular automaton that is induced by the probabilistic skeleton in each mini-column (Fig. 6b) is dead or alive. Fig. 6d shows an example of a wandering activity pattern that typically arises in this cellular automaton. In fact, also very complex periodic and transient activity patterns, reminiscent of dynamic activity patterns in the neocortex (see e.g. Han, Caporale, and Dan, 2008), are known to arise in this particular cellular automaton (Rendell, 2011) for a suitable external network input at the during step 1.

**Figure 6:**
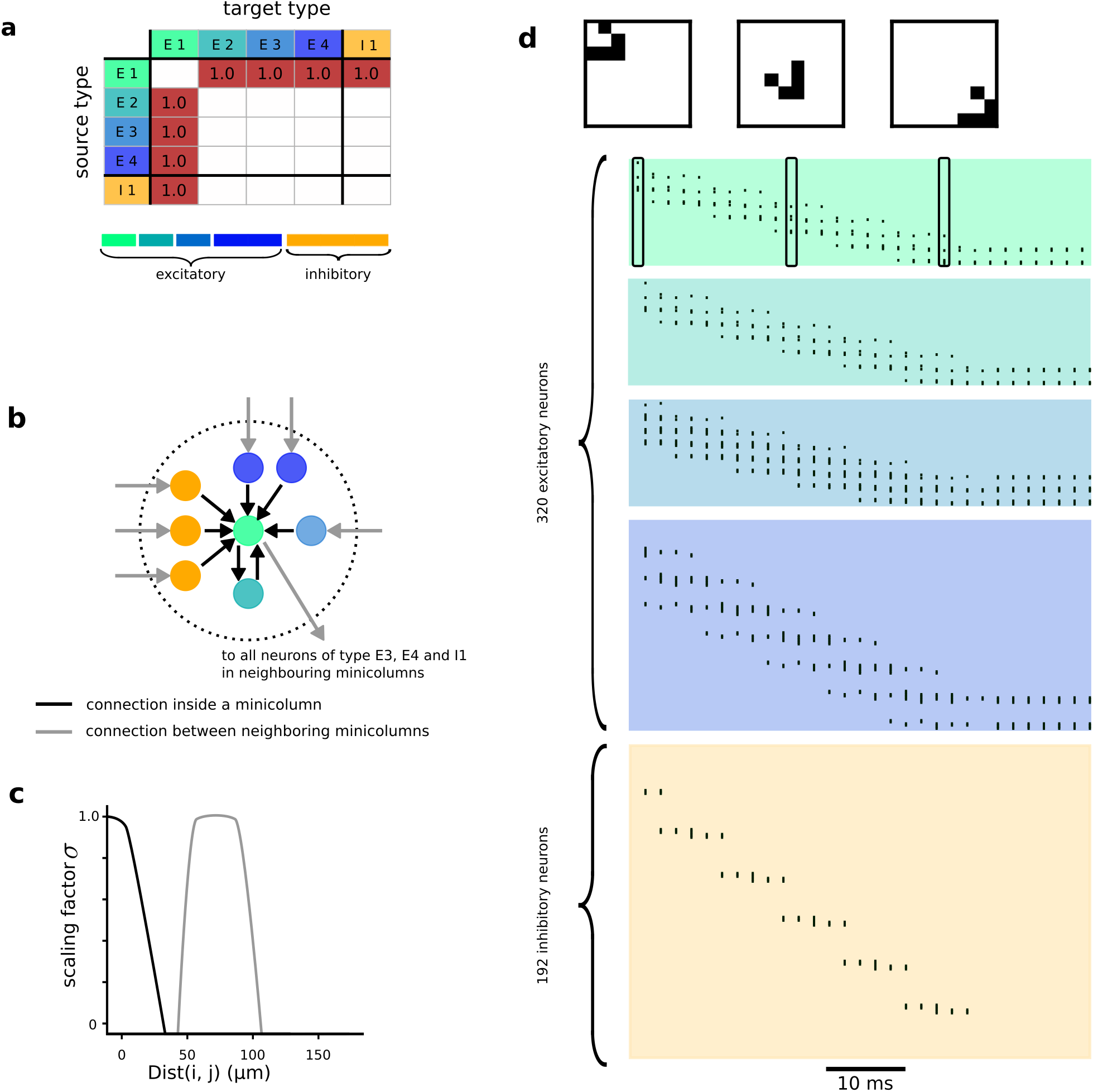
Generation of a spike-based cellular automaton, the Game-of-Life, through a probabilistic skeleton. **a** A probabilistic skeleton that generates the Game-of-Life (only probability values 1 and 0 are needed). **b** Resulting network of spiking neurons within a mini-column. **c** Distance-dependent scaling function for this probabilistic skeleton. The gray curve scales connections from neurons of type E1 to neurons of types E3, E4, I1, the black curve all other connections. **d** Sample traveling activity patterns in an RSNN sample (the patterns shown at the top are encoded by the firing of E1 neurons), and the spiking activity of all neurons in this implementation of the Game-of-Life in a RSNN.

**Figure 7:**
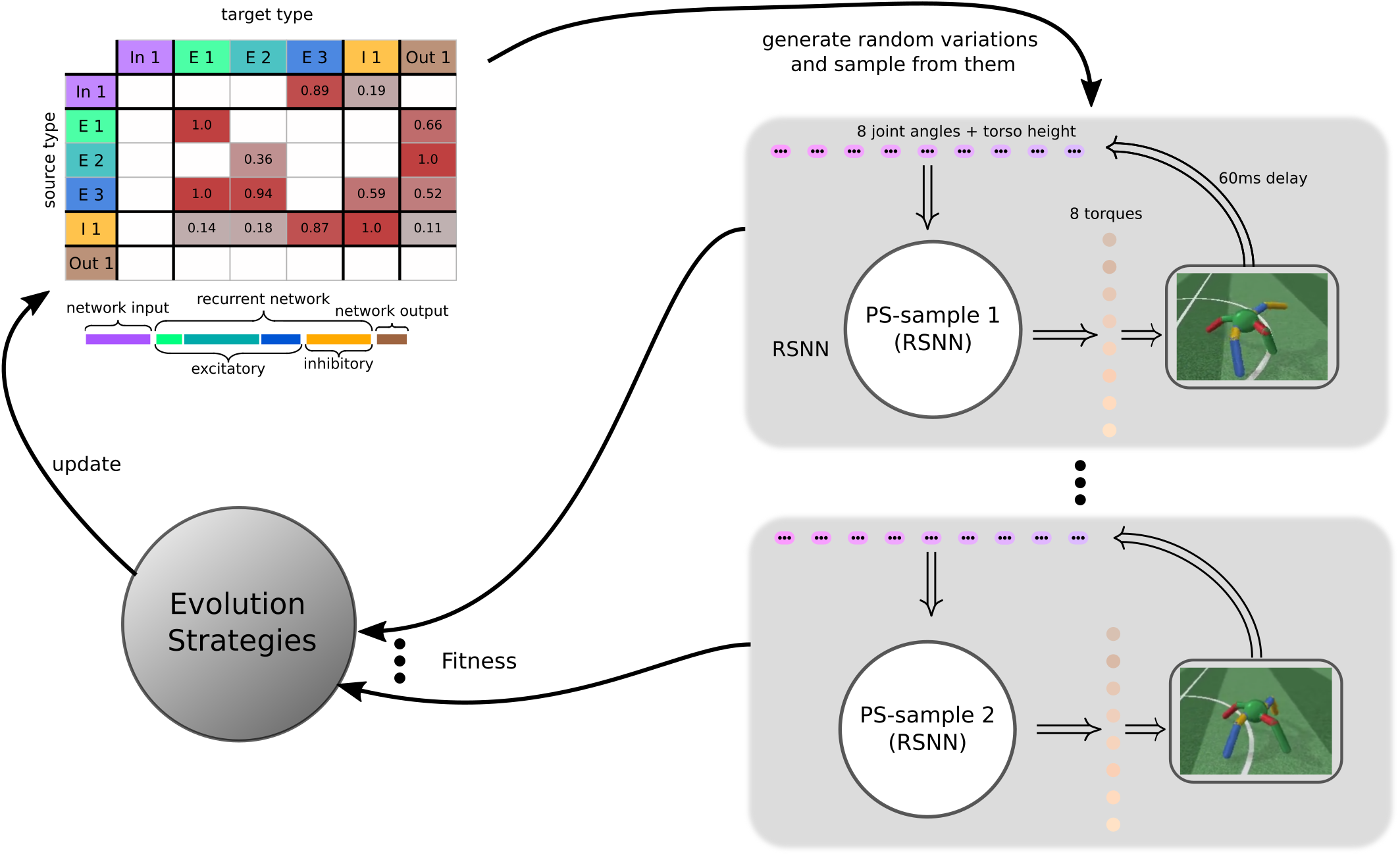
Illustration of our algorithmic approach for optimizing a probabilistic skeleton for a computing task. The motor control task of Fig. 4 is used as illustration. Several RSNNs are sampled from the current probabilistic skeleton, and their capability to solve the given task, i.e., their fitness, is measured. Evolution strategies modify the probabilistic skeleton based on these fitness values. Then the loop is iterated.

Obviously, any finite automaton can be induced in the same way by a special case of a probabilistic skeleton where all connection probabilities have values 0 or 1. Furthermore, also enhanced versions of cellular automata can be induced through a probabilistic skeleton if one allows distance-dependent scaling of connection probabilities that covers a wider range than the simple one needed for the Game-of-Life (Fig. 6c). In particular, also cellular automata that are able to carry out image segmentation through an efficient parallel computation (Sandler et al., 2020), or which can classify external input patterns in a highly parallel manner through intercommunication between cells (Randazzo et al., 2020), can be induced by probabilistic skeletons. Probabilistic skeletons whose connection probabilities are not constrained to the extreme values 0 and 1 are able to induce stochastic versions of such cellular automata that may have additional computational properties such as the capability to solve constraint satisfaction problems, see e.g. Habenschuss, Jonke, and Maass, 2013.

## 3 Discussion

We have addressed two key open problems in theoretical neuroscience: How the genetic code is able to induce complex innate computing capabilities in neural networks of the brain with a small number of parameters, and why the brain employs a fair number of genetically different neuron types. We have shown that distance-modulated neurontype specific connection probabilities between neurons, as found in experimental data on the anatomy of cortical microcircuits Billeh et al., 2020, suffice to induce innate complex computational functions in networks of spiking neurons, provided there exists a substantial number of different neuron types. We have demonstrated that this holds even for the simplest case where neurons of different (genetic) types have the same interal dynamics. The concept of a probabilistic skeleton turned out to be useful for providing this insight, since it encapsulates a known fragment of the programming language which is available to the genetic code for determining the structure of neural circuits. We have shown that probabilistic skeletons are able to induce generic 2D computational operations in cortical microcircuits through their structure. They also allowed us to show that innate pattern recognition capabilities, such as recognition of odors from poisonous food, and innate motor control capabilities can be induced through genetically encoded structure of neural circuits. Surprisingly, also fundamental capabilities to compute with spiking neurons on temporal patterns can be induced through the network structure.

Probabilistic skeletons suggest a particular method for meeting the challenge of Zador, 2019: Understanding the functional impact of the “genomic bottleneck”, i.e., of the fact that the number of bits which the genome uses for encoding neural networks in the brain is really small in comparison with the number of their synapses. A quite different response to this challenge has been addressed by (Koulakov, Shuvaev, and Zador, 2021) on a more abstract level, based on the assumption that the existence of a synaptic connection between two neurons can be determined by linear operations on binary codes for these neurons. The model of (Barabási and Beynon, 2021) is less abstract, and assumes instead that connections between neurons can be formulated as compatibility rules in terms of transcription factors. Implications of the genomic bottleneck on the functional level was demonstrated in these approaches in terms of enhanced generalization capabilities of trained feedforward artificial neural networks.

Our analyses suggests that one should view the neocortex not just as a special case of a deep neural network (DNNs) that acquires its sophisticated computing capabilities, starting with a randomly structured configuration, through supervised gradient descent learning, like DNNs in AI. Rather the cortex can better be captured by computational models that merge aspects of DNNs with aspects of cellular automata (CA), a common model for explaining the emergence of function through structure in 2D arrays of repeating stereotypical “cells”. We have shown that having a fair number of different neuron types enables the genetic code to encode through connection probabilities between different neuron types the computational function of finite automata into neural circuits. Cellular automata are therefore special cases of 2D sheets of neural circuits that can be induced through a probabilistic skeleton, and therefore in principle through the genetic code. Hence probabilistic skeletons, and more generally the principle to encode neural circuits through connection probabilities between different types of neurons, creates a link between RSNNs and cellular automata as dual paradigms for the organization of computational function in the neocortex. Since RSNNs that are generated by a probabilistic skeleton are, unlike cellular automata, not constrained to have only synaptic connections between neurons within the same or in neighboring “cells”, they represent more powerful computational models, especially for fast parallel computation. It should also be noted here that according to experimental data there are also numerous long-range connections in the neocortex, especially between different cortical areas, that are likely to give rise to advanced versions of probabilistic skeletons and RSNN samples with additional innate computing and fast learning capabilities.

Neural networks that are derived as samples of a probabilistic skeletons differ in another aspect from commonly considered network architectures: Their number of synapses and total wire length grows just linearly with the number of neurons. This property is obviously essential for any physical implementation of neural network connections, both in brains and in neuromorphic hardware. In addition, their resilience to weight perturbations supports an implementation of synapses through memristors. Another possible technological application of probabilistic skeletons arises in the domain of organoids (Bhaduri et al., 2020), where it is highly desirable to induce computational function in brain-like organoids through their genetically controlled structure, without invoking synaptic plasticity. Their style of indirect encoding by using different neuron types is also likely to enhance already existing indirect coding approaches in the area of neuroevolution (Ha, A. Dai, and Le, 2016; Stanley et al., 2019). On a more general level, the result that network structure acquires a substantially stronger impact on network function if the network units consist of a fair number of different types suggests a new research direction in network science.

## 4 Methods

### Neuron types

There are 3 kinds of neuron types: input types, recurrent types, and output types. The neurons from these three categories are referred to as input neurons, recurrent neurons, and output neurons.

Input neurons provide external inputs in the form of spike trains. They have no internal states, and there are no recurrent connections from recurrent neurons or output neurons back to the input neurons. The output neurons receive their input from the recurrent neurons (see Fig. 1e).

Recurrent neurons can have connections from input neurons and other recurrent neurons. Each recurrent neuron can only give rise to excitatory neurons or only of inhibitory neurons. Note that input or output types only consist of excitatory neurons.

### Neuron and synapse models

Recurrent and output neurons are modelled as discrete-time versions of standard Leaky-Integrate-and-Fire (LIF) neuron models, More precisely of the GLIF_1_ model from (Teeter et al., 2018). The definition of the continuous neuron model, on which the discrete-time model is based on, can be found in the Suppl. in section 1.3. Control experiments with the GLIF3 model from (Billeh et al., 2020) produced qualitatively similar results.

For the discrete time version of neuron *j* ∈ {1,..., *N*} of type *J* the membrane potential is denoted by *V_j_* and the input current by *I_j_*. We assume that currents are constant on small intervals [*t,t+δt*], which have been set to a length of 1 ms. The neural dynamics of the model in discrete time can then be given as

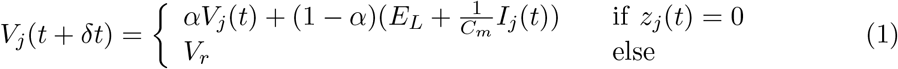

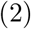

where 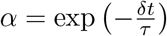 and

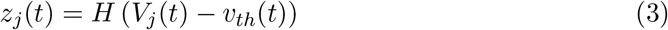

with the Heaviside function 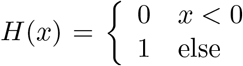. Here *τ* ∈ ℝ is the membrane time constant, *E_L_* ∈ ℝ is the resting potential, *C_m_* ∈ ℝ is the membrane conductance and *v_th_* is the threshold voltage. After spiking the neuron enters a refractory period, lasting *t_ref_* > 0, in which *z_j_*(*t*) is fixed to zero.

The previously defined neuron model use the following set of parameters:

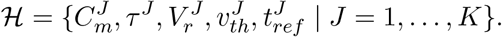

The values for 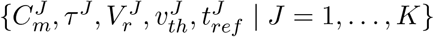 are taken from (Billeh et al., 2020), and the raw data is available in (*V1 Network Models from the Allen Institute* n.d.). A good overview of these neuron types has been made available online in the database of the Allen institute. Detailed biological and modelling data for the prototype of the excitatory neuron can be found at Excitatory neuron^3^ and the prototype for the inhibitory neuron at Inhibitory neuron^4^. We have seen no evidence that the exact values of the GLIF_1_ parameters are essential for the results reported in this paper.

The same synapse model as in (Billeh et al., 2020) has been used. Additional information about the synapse model as well as a mathematically more precise description can be found in the Suppl. in section 1.4 and in Fig. S4.

### Details to the definition of a probabilistic skeleton

A probabilistic skeleton consists of

i. A natural number *K* (the number of neuron types in the model; we have set *K* = 6 in the illustrations of the model in Fig. 1c.
ii. Base connection probabilities *ρ_I→J_* for neurons of type *I* to neurons of type *J*, for the case that they are located within the same minicolumn (see upper part of Fig. 1c for a sample table of such base connection probabilities).
iii. The prevalence *p_I_* of each neuron type *I*, i.e., a number representing the fraction of neurons belonging to type *I* in a generic minicolumn, see the bottom plot of Fig. 1c. Further details can be found in the Suppl., section 1.5.
iv. The common weight *w_in_* of all synapses from input neurons, as well as the common weight *w_E_* of all synapses from excitatory and the common weight *w_I_* of all synapses from inhibitory neurons in the recurrent network.
v. A scaling parameter *σ* that controls the decay of connection probabilities with the horizontal distance between somata.

A probabilistic skeleton is a generative model, which defines a distribution over neural networks of different sizes and with different synaptic connections that share common architectural features.

One samples a neural network from a probabilistic skeleton according to the following rules:

1. Pick a number *n_mcol_* of minicolumns and a number *M* ≥ *K* of neurons per minicolumn. This determines the number of neurons *N* = *n_mcol_* · *M* in the sample network.
2. Draw *S* times for any pair (*i,j*) of neurons with *i* of type *I* and *j* of type *J* from the binomial distribution with probability:

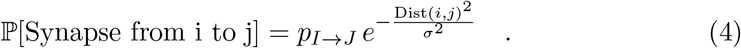 This yields the number *m_ij_* of synaptic connections from *i* to *j*.

The functional form of the dependence of connection probabilities on Dist(*i, j*) approximates the corresponding data from (Billeh et al., 2020), see panels b and d in Fig. 1. We have set *S* = 8 in all our experiments, thereby allowing up to 8 synaptic connections between any pair of neurons. According to Fig. 7A in (Markram et al., 2015) most synaptically connected neurons do in fact have multiple synaptic connections. The effective strength (weight) of a synaptic connection from neuron *i* to neuron *j* is then the product of the general scaling parameter *w_in_, w_E_*, or *w_I_*, that depends on the type of neuron *i*, and the number *m_ij_* of synaptic connections from *i* to *j* that results from drawing *S* = 8 times from the distribution given in equ. (4).

### Optimization method

Probabilistic skeletons were optimized for specific computing tasks with the Separable Natural Evolution Strategy (Separable NES), which had been introduced in (Schaul, Glasmachers, and Schmidhuber, 2011). The algorithm is given below in pseudo code. For the optimization of the *d*-dimensional vector ***θ*** of parameters of the probabilistic skeleton the algorithm uses a Gaussian distribution in every dimension, with means ***μ*** ∈ and variances ***σ*** ∈ ℝ^*d*^. The basic idea is that one samples λ times from this distributions, then evaluates the fitness values of the so-called offsprings, i.e. the vectors 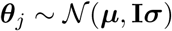, and finally adapts the Gaussian distributions to capture more of those parts of the parameter space where the fitness of the offsprings is higher. The fitness function *F* depends on the computational task for which the probabilistic skeleton is optimized. The mean values of the parameters are initialized by truncated normal random variables with mean zero and variance 1.0 and the variance values are initialized as ones. We found that choosing the learning rate for ***μ*** as *η_μ_* = 1.0 yields good results, which is consistent with the suggested value in (Wierstra et al., 2008) and (Salimans et al., 2017). The learning rate for ***σ*** was chosen as *η_σ_* = 0.01. As suggested in (Salimans et al., 2017) mirrored sampling has been employed, see, e.g., (Brockhoff et al., 2010). That is, for every Gaussian noise vector **s** ∈ ℝ^*d*^ also the offspring, which results from using –**s**, will be evaluated.

#### Algorithm 1 Separable NES

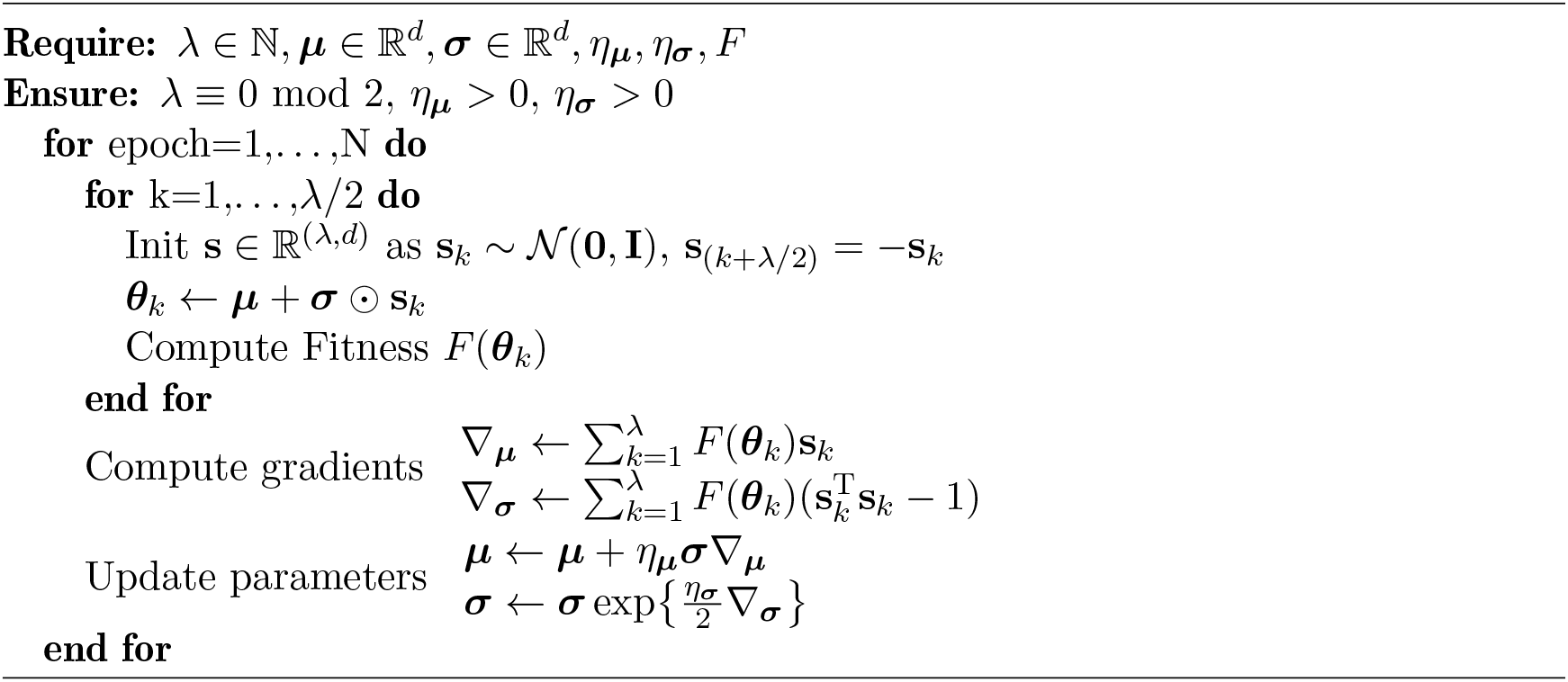

For the optimization of the base connection probabilities *p_I→J_* one needs to make sure that they are always assigned values in [0,1]. For that purpose real valued auxiliary parameters *κ_IJ_* ∈ ℝ are optimized, from which the base connection probabilities are obtained by using the sigmoid function:

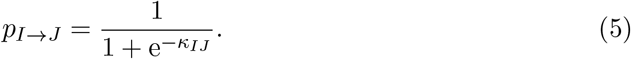

The value of the number *K* of neuron types and of the scaling parameter *σ* from equation (4) were optimized through a separate hyperparameter search.

### Experiments

#### Details to the delayed pattern matching task

##### Task description

In this task two 2D patterns are presented to a RSNN with a variable delay. The goal of the task is for the network to decide whether the two patterns are similar or different. Similar in this context means that the patterns have been sampled from the similar pattern probability distribution, while different would indicate that the patterns originated from two different probability distributions.

###### Input generation

The input generation process is visualized in Fig. 8. To generate a pattern probability distribution a random point with coordinates 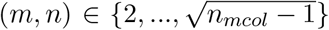 is drawn randomly. Subsequently, a value of 1 is assigned at the coordinates of this point, while all other points have a value of 0, as can be seen in the first part of Fig. 8. This point represents the center of activity. Next, the pattern is convolved with a 2D Gaussian kernel, and scaled to obtain firing probabilities, where the highest probability amounts to 0.2. These firing probabilties are associated with neurons of the input type and can be used to sample input patterns, where spikes are drawn independently for every millisecond. Two pattern probability distributions are considered similar if the centers of activity have a distance of less than 2, while dissimilar pattern probability distributions have a greater distance between the centers of activity.

**Figure 8:**
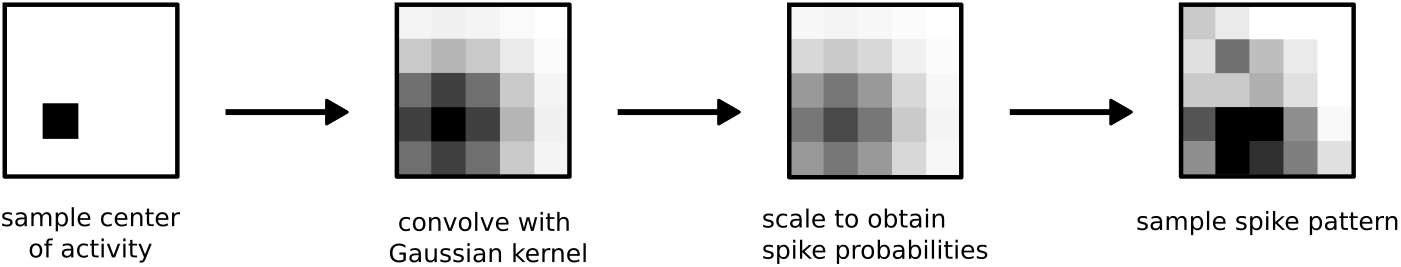
Visualization of the input pattern generation process. In the first step a random center of activity is drawn uniformly on a 2D sheet. Subsequently, a convolution with a Gaussian kernel is applied and the resulting values are scaled and interpreted as firing probabilities. Lastly, these firing probabilities are associated to input neurons and used to sample a spike pattern.

##### Performance measure

The performance measure for this task is the classification accuracy.

##### Fitness function

As a fitness function cross-entropy was used. Furthermore rate regularization was employed to keep the RSNNs from moving to biologically unrealistic firing regimes.

##### Details of probabilistic skeleton and its optimization process

A decay constant of *σ* = 80 was used for this task. The scaling parameters were *w_in_* = 8.66, *w_E_* = 17.6 and *w_I_* = 8.22. The 275 neurons of the RSNN were arranged in *n_mcol_* = 25 minicolumns on a 5×5 grid, where *M* = 11. In every minicolumn there was one input and one output neuron. The probabilistic skeleton contains 74 parameters, whereas the full RSNN contains 61, 875 synaptic weights.

### Details to computations on spike times

#### Task description

The goal is here to classify the temporal distance between two waves of input spikes. There is a fixed time interval of 200 ms, which is divided into four bins of 50 ms. For each class the first spike occurs at the beginning *t* = 0 and the second spike is uniformly drawn from the four bins, which results in four classes of input spike trains. The precise timing of the second spike is again uniformly sampled within the time interval of the chosen bin.

#### Input

The network receives as input a wave of spikes at the beginning of a 200ms long trial, and the second wave at any other time during the trial. For each input neuron some Gaussian noise with mean zero and variance 1 ms has been added to the spike times to avoid that all input neurons spike at the exact same time.

#### Performance measure

The percentage of correctly classified distances between the two waves of input spikes is used as a performance measure. The standard deviation of the performance on this and the subsequent tasks was obtained by averaging over the performance of 50 different RSNNs sampled from the same probabilistic skeleton evaluated on 100 inputs each.

#### Fitness function

Best optimization results are achieved when a different fitness measure than accuracy is used. To compute the fitness the softmax function was applied to the vector (*r*_1_,*r*_2_, *r*_3_, *r*_4_)^*T*^ of spike counts of the 4 output neuron types during the last 30ms of a trial to obtain the class probabilities (*p*_1_,*p*_2_,*p*_3_,*p*_4_)^*T*^. To compute the fitness the target class *y* was first one-hot encoded to the target class vector **y**, i.e. to a vector where all entries are 0 except the element at position *y* – 1, which has the value 1. An example of one-hot encoding can be found in the Suppl., section 1.6. The fitness function is given by the negative cross entropy loss. For a single example with one-hot-encoded target class **y** the fitness is defined as:

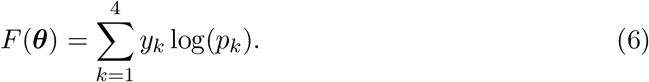

#### Details of the probabilistic skeleton and its optimization process

A decay constant of *σ* = 77.7 was used for this task. The scaling parameters for synaptic strengths were *W_in_* = 14.6, *w_E_* = 15.49, *w_I_* = 6.92. The 304 neurons of RSNN samples during optimization were arranged in *n_mcol_* = 16 minicolumns on a 4×4 grid, where *M* = 19. In every minicolumn there are two input neurons and there is one output neuron per type.

During the optimization of the probabilistic skeleton the activity of output neurons was not only considered during the last 30ms. Instead, initially all spikes of output neurons were counted during the full 200ms of a trial. In the course of the optimization this period was gradually reduced to the last 30ms.

### Details to installing in RSNNs the capability to recognize specific spike patterns

#### Generation of spike inputs

Two clearly distinct ensembles of Poisson spike trains from 4 neurons with a rate of 50 Hz were frozen as templates. Spike input patterns of classes 1 and 2 were generated by creating variations of these spike templates: For every input neuron two time steps from the first 50ms were chosen, and a new spike was inserted at them or the spike was deleted if there was a spike at this time step. Subsequently, the spike times of all spikes in the template were shifted by a random amount drawn from a Gaussian with mean zero and variance 0.5 ms and rounded to the nearest integer value. The third class consisted of random Poisson spike trains over 50ms with a rate of 50 Hz.

#### Input

The network received as input a spike pattern of 4 input neurons over 50ms from one of the three classes, drawn with uniform probability from the three classes.

#### Performance Measure

The same performance measure as for the preceding task was used.

#### Fitness function

A corresponding fitness function as for the preceding task was used.

#### Details of the probabilistic skeleton and its optimization process

Parameters *w_in_* = 14.38, *w_E_* = 7.85, *w_I_* = 7.90 and σ = 129.73 were used. RSNN samples that were tested during the optimization of the probabilistic skeleton consisted of 148 neurons, which were arranged in a 3×4 grid of *n_mcol_* = 12 minicolumns, each minicolumn consisting of *M* = 12 neurons. There was one input neuron in every corner of the grid, hence the corresponding columns had one neuron more than *M*.

### Details to innate motor control capabilities through probabilistic skeletons

#### Task description

For the simulation of the environment (AntMuJoCoEnv-v0) the PyBullet physics engine (Coumans and Bai, 2016–2021) was used. The agent is a quadruped walker and is usually referred to as ‘ant’ in the literature. It consists of four legs with four joints, which are attached by another four joints to a torso, modelled as a sphere. The center of the sphere defines the location of the plant on a 2D plane. The goal of this task is to achieve a high movement speed over the whole trial period, while also avoiding to touch the ground. An episode is terminated if the center of its torso moves below a height of 0.2m, or if the maximum number of time steps has been reached.

#### Spatial structure of RSNN samples

The population coding of continuous-valued input variables induced a prominent 1D dynamics in populations of input neurons, and there seems to be no natural way to map these 2D input arrays properly into a 2D structured RSNN for computational processing. For this reason, and because such basic motor control capabilities are likely to be encoded in the spinal cord and other subcortical structures, we used for this task a 1D arrangement of neurons in order to define their spatial distances, rather than neocortical minicolumns. More precisely, the neurons of input and recurrent types were evenly-spaced distributed over a 1D line segment [0,660] μm. The locations of output neurons, organized for each output variable into two output types consisting of 4 neurons at the same location (see below), were optimized alongside the other parameters of the probabilistic skeleton. The distance measure Dist(*i,j*) for neurons *i* and *j* was computed as the absolute value of the difference between their 1D coordinates.

#### Input

Time in the simulated environment was discretized to time steps of 17 ms length. For this reason, the network received each continuous-valued input value for 17 ms through population coding in one of the 9 input neuron types, each having 16 neurons. It should be noted that population coding is commonly employed in the brain to encode continuous-valued variables (Georgopoulos, Schwartz, and Kettner, 1986). The input was provided by the current state of the simulated environment. Its state space was 111 dimensional. We excluded most of them, for example angular velocities, to have a more compact and arguably biologically more realistic network input.

#### Output

The action space of the controller is given by 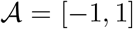^8^, which corresponds to 8 torques applied to the 8 joints of the ant. An output torque *y* ∈ [–1, 1] of the model is computed by using two output neuron types, each consisting of 4 output neurons, representing negative and positive torques to a joint, denoted by *J*_−_ and *J*_+_. This corresponds to motor commands in the form of firing rates to 2 antagonistic muscles for a joint. Firing activity of output neurons of the RSNN were decoded as signal to the simulated environment by computing the normalized linear combination of the spike rates over a 17 ms time step of the environment:

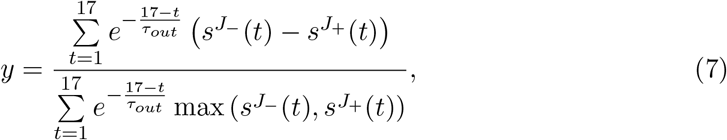

where *τ_out_* = 10.

#### Performance measure and fitness function

The performance measure was the same as the fitness value. The fitness was given by the total reward received from the environment, summed up over time. At every time step of 17ms length the agent received a reward

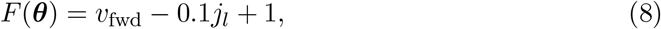

where *v*_fwd_ is the velocity of the center of the ant in the *x* direction, *j_l_*: = number of joints which are at the limit. A constant reward of 1 was added for each time step in order to induce long lasting locomotion without premature abortion of an episode because the torso touched the ground.

RSNN samples with 458 neurons from the optimized probabilistic skeleton produced an average fitness of 517 (standard deviation of 51.85) using 250 steps in the environment, where the average was computed over 100 trials. The version of the model where 30% of the recurrent neurons are randomly deleted achieved an average fitness of 331.

#### Details of the probabilistic skeleton

The probabilistic skeleton consisted of *K* = 40 types, and was optimized for RSNN samples with *N* = 458 neurons. Every input type was constrained to only form connections to one recurrent type, which did not receive synaptic inputs from another input type. The other parameters were *w_in_* = 4.75, *w_E_* = 4.5, *w_I_* = 2.3 and *σ* = 80.0.

Note: The version of the ant locomotion task that we considered differed somewhat from the version that is commonly considered in the literature (Schulman et al., 2015). There one does not assume a delay in feedback from the environment. Also, the more limited observation space that we used made it harder for the model to know in which direction it was facing, especially at a later point in the trial. This made it harder to move especially along the x-axis, which was the only direction in which locomotion was rewarded.

### Details to game of life

#### Task description

The goal of this task is to demonstrate that a PS can generate an arbitrarily large RSNN which is capable of simulating the cellular automata game of life.

#### Input generation

There is no input for this task. One could consider the initial state of the recurrent network to be the input.

#### Performance measure

There is no performance measure for this task.

#### Fitness function

There is no fitness function for this task.

#### Details of the probabilistic skeleton and its optimization process

For this task a different paradigm for spatially dependent probability scaling has been considered, see Fig. 6c. The scaling parameters were *w_E_* = 1 and *w_I_* = 1. There are no input and output types for this task. The baseline connection probabilities *p_I→J_* have not been optimized using ES. Instead they have been computed analytically. Note, that for this task very simple McCulloch-Pitts neurons have been considered. In theory, game of life should be played on an infinitely large cellular automata, but as this would require an infinite amount of resources to simulate. As our simulations only use a finite cellular automata the behavior at the boundaries can diverge from what would be expected from an infinite field.

### Details to Figure 5

To compare the different tasks it is necessary to use for different computing tasks a common performance scale. This can be achieved by defining the baseline for every task as the performance level of a random output. For example, the computations on spike times task required a decision between 4 classes, hence picking a random class would give for a uniform distribution of classes an expected accuracy of 25%. Analogously the baseline accuracy for the spike pattern classification, which involves three classes, is 33.33%. The baseline for the ant was considered to be a reward of 150, which amounts to the reward received after 250 time steps without moving forward. For the highest performance was defined by the performance of the best probabilistic skeleton.

The performances on these different tasks were scaled by calculating for each task the difference between the theoretically best possible performance (either accuracy or normalized cross correlation) to the baseline performance and normalizing this difference to [0, 1].

For panel a, each number of recurrent neuron types 80 probabilistic skeletons were optimized for every task, and the best performing ones were used for the plot.

For panel b, the effective weight of each individual synapse was independently perturbed for each presynaptic spike. The amplitude of this perturbation was measured as fraction x of its current value, and the maximal fraction is indicated on the x-axis of the panel. For each value of x the noise value was drawn uniformly from the interval [—x,x]. The resulting perturbed weight was set to zero if the perturbation caused its sign to change.

#### Details to the comparison of neuron density, synapses numbers, and wire length with experimental data from the neocortex

According to Fig. 2B of (Carlo and Stevens, 2013) the number of neurons under a square mm of the neocortical sheet is in the mammalian brain around 100,000. The number of synapses per neuron was estimated in (Braitenberg and Schüz, 2013) to be 7777, and the total length of axons per neuron was estimated to be 4.4cm. We have compared these experimental data with corresponding estimates that arise for RSNN samples from probabilistic skeletons for the computing tasks that we considered (see Table 1 in the Suppl.). For example, the RSNN for coincidence detection, whose firing activity and performance was shown in Fig. S2 f-h, has 2160 neurons, occupies a square patch of 0.5184*mm*^2^, has 360,100 synapses, and a total wire length of 17.5m. Thus its number of neurons per square mm is by a factor 22 smaller than in the mammalian brain, the number of synapses is by a factor 1008 smaller, and its total wire length is by a factor 118 smaller than in the data. Thus, these numbers are in a reasonable range, but significantly smaller than in the experimental data. The main reason for that is that the number of neuron types that are needed for each of the computing tasks that we considered is substantially smaller than the estimated 111 neuron types in mouse V1 (Tasic et al., 2018). Consistent with that, the number M of neurons in a minicolumn was in our examples well below the 80 -,120 neurons in a typical neocortical minicolumn. Note that the number of synapses and total wire length grow superlinearly with the number of neuron types, (see Suppl. section 1.7 and 1.8). In addition, we only counted wire length in the horizontal direction, and ignored long-range connections.

## Acknowledgements

We would like to thank Dániel Barabasi, Guozhang Chen, Peter Jonas, Eben Kadile, Robert Legenstein, Jason MacLean, Risto Miikkulainen, Franz Scherr, Kenneth Stanley, and Yuqing Zhu for helpful comments on a prior version of this manuscript. This research was partially supported by the Human Brain Project (Grant Agreement number 785907) of the European Union. Computations were carried out on the Human Brain Project PCP Pilot Systems at the Juelich Supercomputing Centre, which received cofunding from the European Union (Grant Agreement number 604102) and on the Vienna Scientific Cluster (VSC).

## Author contributions

WM and CS designed the approach, CS and DL carried out the experiments and analyzed the results, WM, CS and DL wrote the paper.

## Footnotes

1 https://cloud.tugraz.at/index.php/s/iXDSo6Q7HDmDyX6

2 https://cloud.tugraz.at/index.php/s/WpyRncz62p9PnTc

3 https://celltypes.brain-map.org/experiment/electrophysiology/501848315

4 https://celltypes.brain-map.org/experiment/electrophysiology/313862167

