## Supplements for "Structure induces computational function in networks with diverse types of spiking neurons"

**2 Supplementary figures** **17**

### 1 Supplementary notes

#### 1.1 Connectivity diagram of the probabilistic skeleton for the pattern change detection task

The chord diagram in Fig. S1 is drawn in the style of Fig. 7C of (Markram et al. 2015), using the base connection probabilities $p_{I \to J}$. The thickness of each chord reflects the value of the corresponding base connection probability and the length of each segment (arc) on the perimeter is proportional to the expected fraction of synapses from and to the corresponding neuron type. Such a chord graph of a probabilistic skeleton should not be confused with the connectivity graphs of individual RSNN samples from this probabilistic skeleton. The latter also depends on the selected number $N$ of neurons, the value of $\sigma$, and numerous outcomes of random drawings. More precisely, the arc length of type $I$ is proportional to:

$$\frac{\sum_J p_{I \to J} + \sum_J p_{J \to I}}{2 \sum_{IJ} p_{I \to J}}.$$

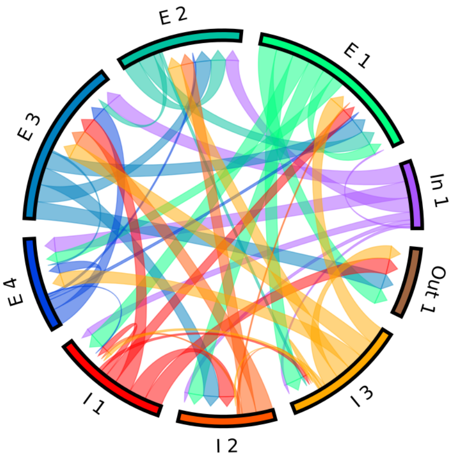

**Figure S1:** Connectivity diagram of the probabilistic skeleton for the pattern change detection task. Connectivity graph induced by this probabilistic skeleton, plotted by a chord diagram similarly as used to report experimental data (see Fig. 7C of Markram et al. 2015)

#### 1.2 Further Generic 2D computing capabilities.

The neocortex forms a 2D sheet, with topographic maps between cortical areas and from peripheral sensory neurons, as well as between layers within a cortical area. It is commonly conjectured that the neocortex is genetically endowed with basic computing capabilities that make use of this 2D organization (Mountcastle 1998). Hence an important test for our theory that functional capabilities are induced in neural networks of the neocortex through their structure, i.e., through a probabilistic skeleton, is whether basic 2D computing capabilities can be induced by them. We demonstrate here for two further tasks, besides the delayed pattern matching task of Fig. 2, that this is in fact the case.

Contrast enhancement and amplification of local maxima, i.e., applying a Mexican hat filter, are further standard examples for such a basic 2D computing capabilities of generic cortical microcircuits. We found that a probabilistic skeleton with just 4 recurrent neuron types (see Fig. S3a) endows RSNN samples with the capability to approximate the computation of a Mexican hat filter, achieving for example for 5x5 input patterns a correlation of 0.89 with this target. Fig. S2c and d depict two examples for the resulting transformation of the input patterns shown at the top to the outputs shown at the bottom by an RSNN sample from the probabilistic skeleton with 1152 neurons. The grey values of pixels in the input- and output patterns were encoded through firing rates. While this RSNN sample had 1152 neurons and 995, 328 potential synaptic connections, the probabilistic skeleton encoded this computation with just 32 parameters. Although this probabilistic skeleton had been optimized for computations on 5x5 inputs (red square in Fig. S2b), RSNN samples could also process the shown 12x12 input patterns very well (cross correlation 0.71 with target). Hence one can argue that the underlying probabilistic skeleton captures a generic 2D computing capability. In particular, the probabilistic skeleton managed to organize the somewhat delicate interaction of excitatory and inhibitory neurons that is needed to execute this task. The phasic firing pattern of the output neurons in Fig. S2a suggests that the resulting RSNN implements the local contrast enhancement through an iterated temporal competition between excitatory neurons in the recurrent network, where the strongest activated neurons fire first, thereby inhibiting weaker competitors -but transiently also themselves.

Cortical areas typically receive synaptic inputs in the form of multiple topographic maps from other cortical areas, and also from the visual or somato-sensory periphery. Coincidence detection between bottom-up and top-down projections is conjectured to be a fundamental operation in perception (Larkum 2013). This computation requires the RSNN to approximate the computation of products of input values at the same location in the two input patterns (see Methods and target outputs at the bottom of Fig. S2f and g), rather than sums: Hence is somewhat nontrivial to implement with spiking neurons. Fig. S2e-g shows that also this fundamental 2D computing capability can be induced through the probabilistic skeleton shown in Fig. S3b. Fig. S2f, g depict the computation of an RSNN sample on two sets of 2D patterns input1 and input2, together with the RSNN output and the target output. Input- and output values were again encoded by firing rates. The RSNN produced the output shown below them. The target output is depicted at the bottom. The RSNN achieved on 5x5 patterns a correlation with the target output of 0.777, which is not optimal, but substantially higher than what is achievable with a linear computation (sum), see Fig. S3c. This probabilistic skeleton had 121 parameters. It had been optimized, like the one for local contrast enhancement, for 5x5 input patterns, but also achieved for 12x12 input patterns a cross-correlation of 0.71 with the target. The RSNNs for the 12x12 inputs had 2160 neurons and 3,732,480 potential synaptic weights. Altogether the results of this section provide evidence that fundamental computational operations that have been conjectured to be employed by cortical microcircuits throughout the neocortical sheet can very well be engraved into these circuits by genetically controlled features of their architecture, in a quite low-dimensional parameter space.

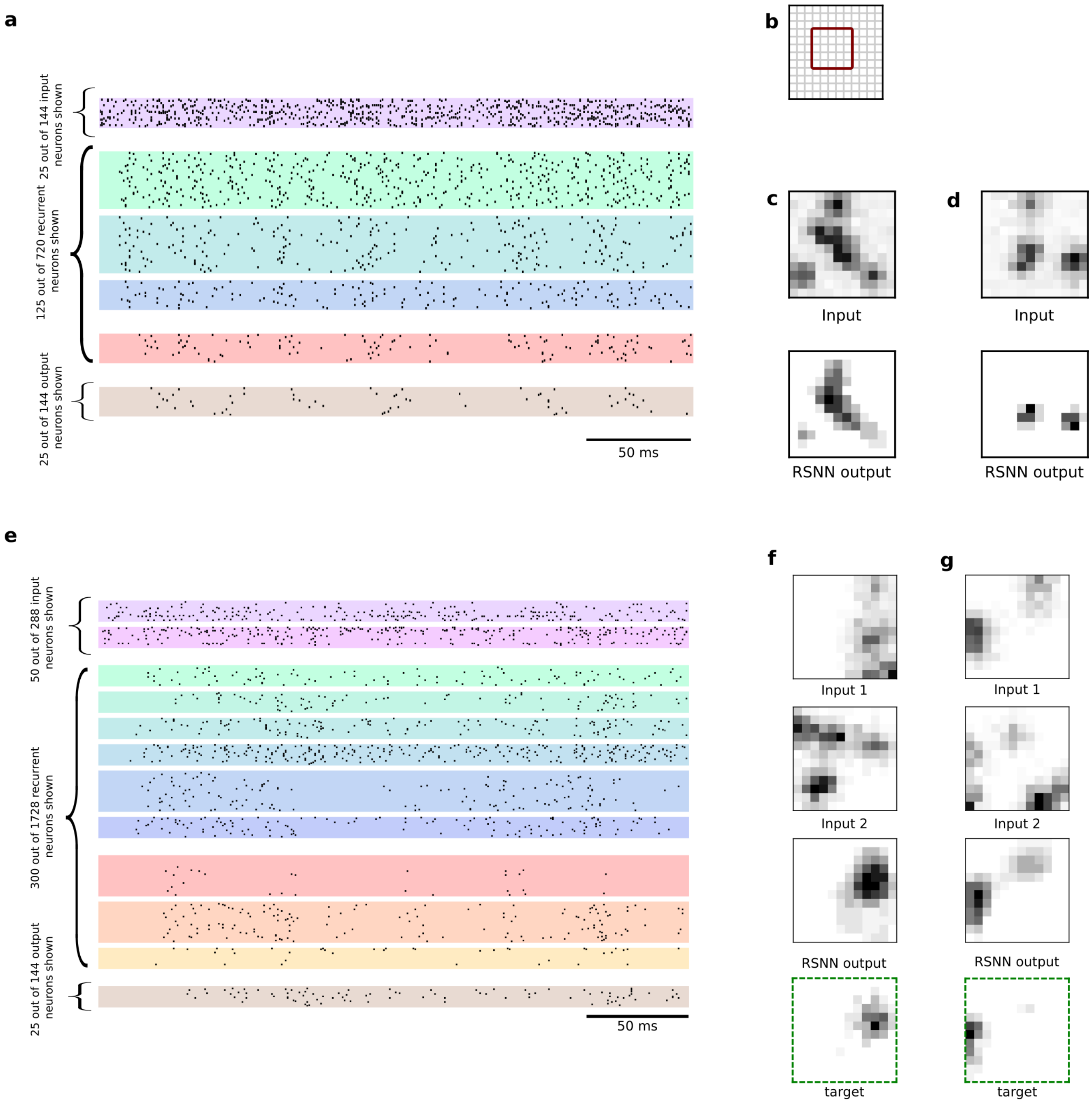
a
25 out of 144 input neurons shown
125 out of 720 recurrent neurons shown
25 out of 144 output neurons shown
50 ms
b
c
Input
RSNN output
d
Input
RSNN output
e
50 out of 288 input neurons shown
300 out of 1728 recurrent neurons shown
25 out of 144 output neurons shown
50 ms
f
Input 1
Input 2
RSNN output
target
g
Input 1
Input 2
RSNN output
target

**Figure S2: Generic 2D computations: Local contrast enhancement and detection of coincidences in two input streams. a, c** Spike raster plot of an RSNN sample solving the local contrast enhancement task, for the 2D input pattern shown at the top of panel c, producing the 2D output shown at the bottom of panel c. Grey values were encoded through Poisson firing rates. **b** The red box in the 12x12 grid indicates the 5x5 pattern size for which the probabilistic skeleton had been optimized, the 12x12 grid the pattern size on which its RSNN sample had been tested (also for the task discussed below). **d** A different 2D input and output pattern for the same RSNN as in c. **e** Spike raster of an RSNN sample of the probabilistic skeleton for solving the 2D coincidence detection task. **f** Pairs of 2D input patterns, for which this RSNN produced the 2D output patterns shown below them. Target output is shown at the bottom. **g** Another sample for the same RSNN.

#### Details to local contrast enhancement

**Task description:** In this task 2D patterns $\mathbf{x} \in [0,1]^{\sqrt{n_{mcol}} \times \sqrt{n_{mcol}}}$ are presented to the network, where $n_{mcol}$ is the number of minicolumns; these are arranged in a square grid. For optimizing the probabilistic skeletons, $n_{mcol} = 25$ was selected, resulting in a 5x5 grid.

**Input generation** To generate a pattern $\mathbf{x}$ first a fixed number $N_{points}$ of coordinate pairs $(m,n) \in \{2,\ldots,\sqrt{n_{mcol}}-1\}^2$ were drawn randomly. In a second step, the value 1.0 was assigned to these points, all other points were set to zero. Finally, a discrete 2D Gaussian filter with variance 1.25 is applied to this binary matrix and the result is normalized to $[c,1]$, where $c$ is a baseline intensity, which is drawn uniformly from $[0.05, 0.15]$. For the 12x12 patterns that can be seen in Fig. S2. $N_{points} = 6$ was used. $N_{points} = 2$ was used for 5x5 input patterns. The resulting 2D patterns were presented to the RSNN through corresponding 2D arrays of Poisson spike trains. Their rates between 0 and 200 Hz were linearly scaled by the grey-values of the corresponding grid cells of the 2D pattern. The length of a trial was 300 ms.

**Targets for the computation:** Target values were computed by applying the Mexican hat filter defined below, with symmetric padding to the 2D input pattern, followed by clipping at zero and normalization to $[0,1]$. The Mexican hat was defined by:

$$\text{filter} = \begin{bmatrix} 0 & -1 & 0 \\ -1 & 4 & -1 \\ 0 & -1 & 0 \end{bmatrix} \tag{1}$$

**Output:** There were $n_{mcol}$ output neurons, one in each minicolumn, which were arranged in a square grid. The output value of each output neuron $i$ was computed by counting its spikes and normalizing that value:

$$\hat{s}_i = \sum_t z_i(t), \tag{2}$$

$$s_i = \frac{\hat{s}_i}{\max(\hat{s})}. \tag{3}$$

**Performance Measure and fitness function:** Fitness was the normalized cross correlation between target signal $\mathbf{y}$ and output signal $\mathbf{s}$:

$$F(\boldsymbol{\theta}) = NCC = \frac{\sum_{k=1}^{n_{mcol}} s_k y_k}{\sqrt{\sum_{k=1}^{n_{mcol}} s_k^2 \sum_{k=1}^{n_{mcol}} y_k^2}}$$

This value can easily be interpreted: A value of 0.0 would indicate that the two patterns are totally different while a value of 1.0 would show that the two patterns are identical. Hence we used it also for measuring performance.

**Details of the probabilistic skeleton and its optimization process:** The probabilistic skeleton, shown in Fig. S3a, had $K = 4$ types and was optimized for RSNN samples with $n_{mcol} = 25$ minicolumns (arranged in a square grid) and $M = 8$ neurons per column, resulting in $N = 200$ neurons of RSNN samples. Other parameters of the probabilistic skeleton were $w_{in} = 1.70$, $w_E = 1.71$, $w_I = 2.15$, and $\sigma = 73.6$.

##### Details to the coincidence detection

**Task description:** Two 12x12 input patterns were simultaneously presented to the network. The goal was to mark positions in the 12x12 grid where both input patterns were reasonably strong, using a product operation.

**Input:** The input patterns were generated using the same algorithm as for the contrast enhancement task, with the only difference that a minimum distance of 3 was imposed on the points used to generate the input pattern.

**Computing the targets:** The target outputs of the coincidence detection task were computed by the following formula:

$$\mathbf{y} = \frac{\boldsymbol{\phi}(\mathbf{x}_1 \odot \mathbf{x}_2)}{\max(\boldsymbol{\phi}(\mathbf{x}_1 \odot \mathbf{x}_2))} \tag{4}$$

where $\mathbf{x}_1$ and $\mathbf{x}_2$ are the two input patterns. $\odot$ refers to the element wise product, and $\phi$ is a function that returns element wise the identity of the input if the element is higher than the average of all vector coordinates of the input vector, else it returns zero, as described in equation 5:

$$\boldsymbol{\phi}(\mathbf{v}) = \begin{pmatrix} \phi(v_1) \\ \vdots \\ \phi(v_n) \end{pmatrix}, \quad \phi(v_k) = \begin{cases} v_k & v_k \geq \text{mean}(\mathbf{v}) \\ 0 & \text{else.} \end{cases} \tag{5}$$

**Output, performance measure and fitness function:** The same output convention and fitness function, and performance measure as for the contrast enhancement task were used.

**Details of the probabilistic skeleton and its optimization process:** The probabilistic skeleton, shown in Fig. S3b, used $K = 12$ neuron types and $M = 15$ neurons per minicolumn. Its other parameters were $w_{in} = 2.28$, $w_E = 3.08$, $w_I = 1.92$, and $\sigma = 70$.

Note, that in the coincidence detection task, the target output is not the sum of two input values, but their product. This is salient, since simply computing a sum and thresholding its value is a trivial task for an RSNN. However, approximating a product is substantially more challenging. In Fig. S3c it becomes apparent that simply computing the sum would not be a good strategy, as it would yield a low fitness value.

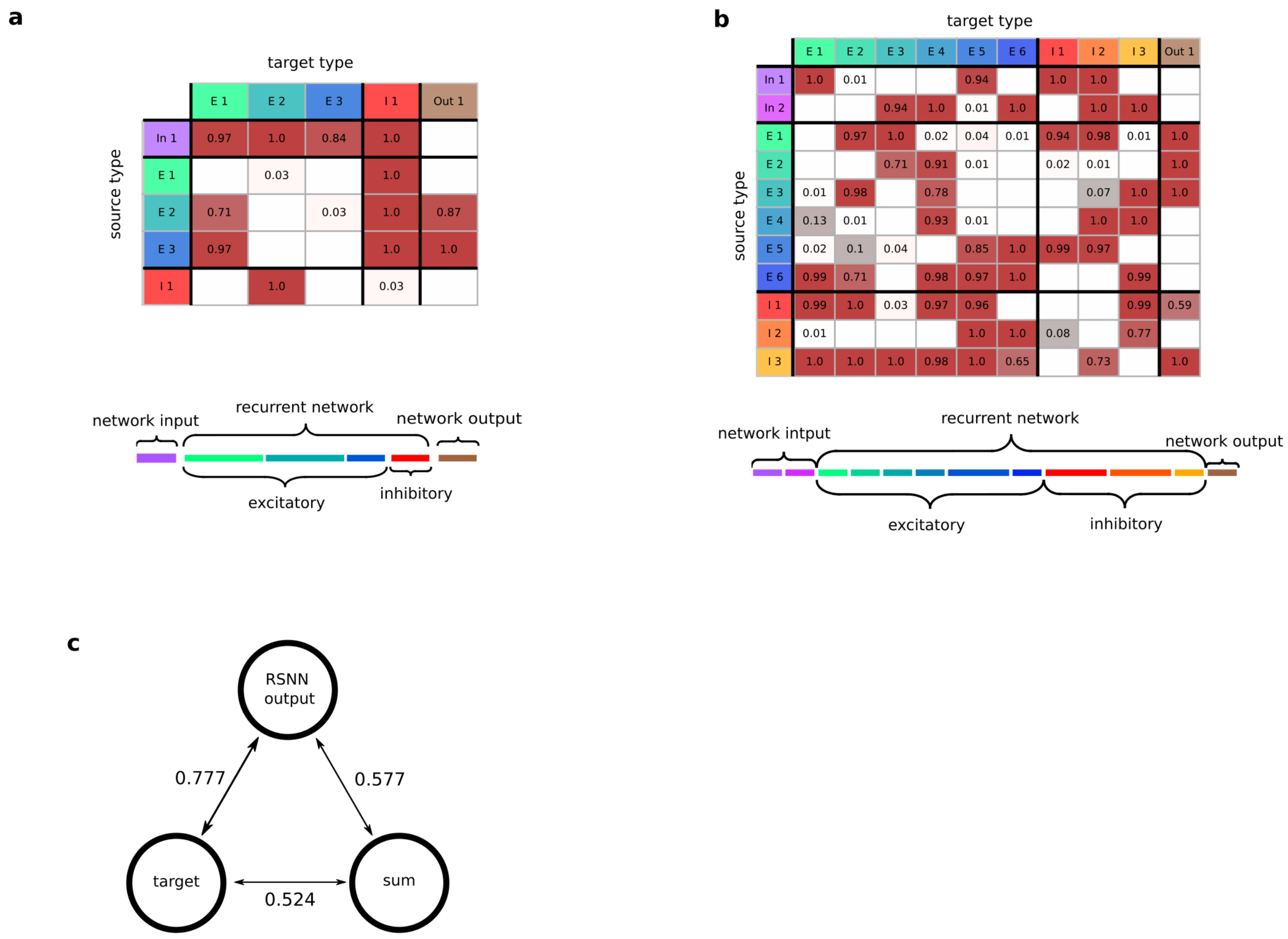

**Figure S3: Details to the induction of 2D computing capabilities through probabilistic skeletons. a** Probabilistic skeleton for local contrast enhancement task. **b** Probabilistic skeleton for the coincidence detection task. **c** Normalized cross entropy between the output of the RSNN for the coincidence detection task, the target and the average of the two input patterns. This shows that approximating the coincidence detection target with a linear operation yields a substantially smaller correlation.

#### 1.3 Continuous neuron model

The continuous neuron model is the conceptual basis for the discrete-time version of the neuron model that is actually used in our simulations. For a given neuron $j \in \{1, \ldots, N\}$ of type $J$ we denote by $V_j : \mathbb{R} \to \mathbb{R}$ the membrane potential and by $I_j : \mathbb{R} \to \mathbb{R}$ the input current. We used different parameters for excitatory and inhibitory neurons that were based on experimental data, see below. Hence these parameters depend for a neuron of type $J$ on whether this is an excitatory or inhibitory neuron type. But for simplicity we will drop in our notation the dependence of the neuron model on the neuron type $J$ until the section below on parameters for neuron and synapse models.

For example, for the membrane conductance we could write $C_m^J \in \mathbb{R}$. Since we will assume for the whole section that we use the same type $J$ we will drop this superscript to simplify the notation, hence

$$C_m = C_m^J.$$

We denote by $\tau \in \mathbb{R}$ the membrane time constant and by $E_L \in \mathbb{R}$ the resting potential. The classic leaky integrate-and-fire linear differential equations reads

$$\tau \frac{d\,V_j(t)}{dt} = -(V_j(t) - E_L) + \frac{1}{C_m} I_j(t). \tag{6}$$

If the voltage is above the threshold $v_{th}$ the parameters get updated by

$$V_j(t+) \leftarrow V_r;\ j = 1, \cdots, N, \tag{7}$$

where $V_r \in \mathbb{R}$ is the reset voltage.

#### 1.4 Synapse model

The time course of a postsynaptic current is modeled like in Billeh et al. 2020 by a linear increase followed by an exponential decay:

$$I_{syn}(t) = \frac{e}{\tau_{syn}} t\,\delta t\, e^{-\frac{t\delta t}{\tau_{syn}}}, \quad t \in \mathbb{N}. \tag{8}$$

$\tau_{syn}$ is the synaptic time constant after which the current amplitude will be at its maximum. Note that $\tau_{syn}$ depends on the types of the pre- and postsynaptic neurons. The exact values of $\tau_{syn}$ have been set to 5.5 ms for excitatory-to-excitatory synapses, 8.5 ms for inhibitory-to-excitatory synapses, 2.8 ms for excitatory-to-inhibitory synapses and 5.8 ms for inhibitory-to-inhibitory synapses, according to Billeh et al. 2020. The current will be negative for synapses from inhibitory neurons. The resulting postsynaptic currents can be seen in Figure S4.

These currents $I_{syn}(t)$ are scaled for each synapse by a general scaling factor $w$ equal to $w_{in}$, $w_E$, or $w_I$, depending on whether the presynaptic neuron is an input neuron, or some other excitatory or inhibitory neuron. Furthermore, they are multiplied for a synaptic connection from neuron $i$ to neuron $j$ by the number $m_{ij}$ of synaptic connections from i to j. Hence, the current from neuron $i$ to neuron $j$ at time $t$ can be written in terms of the spike train $z_i(s)$ of the presynaptic neuron as

$$I_{ij}(t) = \sum_{k=0}^{t-1} \frac{w\ m_{ij}\ z_i(k)}{\tau_{syn}} e(t-k)\,\delta t\, e^{-\frac{(t-k)\delta t}{\tau_{syn}}}. \tag{9}$$

The input current $I_j(t)$ for neuron $j$ at time $t$ is defined as the sum of these currents over all presynaptic neurons $i$.

Note, that there is a transmission delay from the creation of a spike until the arrival at a synapse. For excitatory neurons this transmission delay is 3ms and for inhibitory neurons it is 2ms.

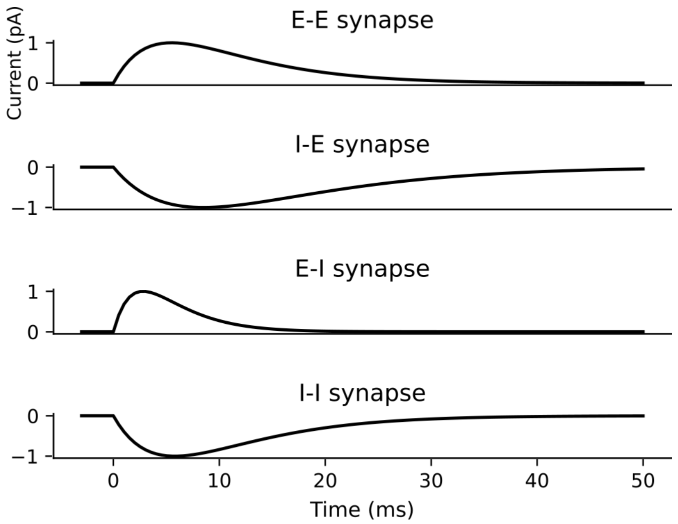

**Figure S4:** The temporal dynamics $I_{syn}(t)$ of the four different synapse types, as used in Billeh et al. 2020.

#### 1.5 Computation of the neuron prevalence

For the optimization of probabilistic skeletons it is convenient to define neuron prevalences as real values: $p_I$ of type $I$ is a real value that correlates to the fraction of neurons which will belong to type $I$ in a generic minicolumn. In order to compute a corresponding assignment of the neurons in a minicolumn to the neuron types we proceed as follows: We apply the softmax function, followed by an adjustment procedure which makes sure that every type has at least one neuron per column and that the total number of neurons stays constant.

The exact algorithm used to obtain the number of neurons per type can be found in algorithm 2.

**Algorithm 2** Algorithm for computing the number of neurons per type using the prevalence $p$, the number of minicolumns $m_{ncol}$ and the total number of recurrent neurons $N_{rec} = (K+3) \cdot n_{mcol}$ (For the contrast enhancement task it is $N_{rec} = (K+2) \cdot n_{mcol}$. The floor function rounds its argument down to the next integer, the maximum function returns the higher valued argument, argsort returns a list of sorted indices, pointing to the highest elements in the argument array first, and the $+=$ operator increments a variable by the amount on the right side.

**Require:** $p$, $N_{rec}$, $n_{mcol}$,
  frac = softmax($p$)
  nnpt = frac $\cdot$ $N_{rec}$
  nnpt = floor(nnpt/$n_{mcol}$)
  nnpt = maximum(nnpt, $n_{mcol}$)
  diff = frac $\cdot$ $N_{rec}$ - nnpt
  max_idx = argsort(diff)
  overlap = floor(($N_{rec}$ - sum(nnpt)) / $n_{mcol}$)
  **for** $k = 1, \ldots,$ overlap **do**
    nnpt[max_idx[k]] += $n_{mcol}$
  **end for**
  **return** nnpt

#### 1.6 One-hot encoding

To convert the target label $y$ to a one-hot encoded vector one first creates a vector with a length equal to the number of classes consisting only of zeros. Then at the position $y-1$ the value 1 is entered. For example, if one considers a setting with four classes and the target class is $y = 3$ this would yield a vector of:

$$\begin{bmatrix} 0 \\ 0 \\ 1 \\ 0 \end{bmatrix} . \tag{10}$$

#### 1.7 Derivation of the expected number of synaptic connections of a single neuron

To derive the expected number of outgoing synapses $ST(i)$ a neuron $i$ can form, we consider an infinitely large grid of minicolums. To obtain the desired estimate one can consider neighboring minicolumns to be located on concentric squares around the source neuron $i$.

Let the function $f(k)$ denote the number of columns in the $k$th concentric square:

$$f(k) = \begin{cases} 1 & \text{if } k = 0 \\ 8k & \text{else} \end{cases} . \tag{11}$$

Then the expected number of outgoing synapses from neuron $i$ can be written as:

$$ST(i) = \sum_{k=0}^{\infty} \sum_{J} f(k) M_J S p_{I\to J} e^{-\left(\frac{\alpha k}{\sigma}\right)^2} \tag{12}$$

, where the first $k$ will sum over all concentric squares, $M_J$ is the number of neurons of type $J$ in a minicolumn, $S = 8$ is the maximum number of synapses between two neurons, $\alpha = 60\mu m$, and $\sigma$ accounts for the distance decay. Note, using $\alpha = 60\mu m$ will yield an upper bound, as the distance to most minicolumns on the concentric square $k$ will be larger than $\alpha k$.

An upper bound of this equation is of interest and can be derived:

$$ST(i) = \sum_{k=0}^{\infty} \sum_{J} f(k) M_J S p_{I\to J} e^{-\left(\frac{\alpha k}{\sigma}\right)^2} \tag{13}$$

$$= S \sum_{J} M_J p_{I\to J} \sum_{k=0}^{\infty} f(k) \gamma^{k^2} \tag{14}$$

, where

$$\gamma = e^{\left(\frac{\alpha}{\sigma}\right)^2}. \tag{15}$$

Breaking up the function $f(k)$ yields:

$$ST(i) = S \sum_{J} M_J p_{I\to J} \left(1 + 8 \sum_{k=1}^{\infty} k \gamma^{k^2}\right). \tag{16}$$

We get an upper bound by using a standard estimate for the geometric series:

$$S \sum_{J} M_J p_{I\to J} \left(1 + 8 \sum_{k=1}^{\infty} k \gamma^{k^2}\right) < S \sum_{J} M_J p_{I\to J} \left(1 + 8 \sum_{k=1}^{\infty} k \gamma^{k}\right) \tag{17}$$

$$= S \sum_{J} M_J p_{I\to J} \left(1 + 8 \frac{\gamma}{(1-\gamma)^2}\right). \tag{18}$$

Note that this upper bound does not contain anymore an infinite sum.

#### 1.8 Expected wire length

An upper bound for the expected wire length can be computed in a similar fashion. The expected wire length per neuron can be written as:

$$WL(i) = \sum_{k=1}^{\infty} f(k) \sum_{J} \alpha k M_J p_{I\to J} \left(1 - \left(1 - e^{-\left(\frac{\alpha k}{\sigma}\right)^2}\right)^S\right) \tag{19}$$

Note, that although there can be up to $S$ synapses between two given neurons, but the wire distance is only counted once, based on the assumption that multiple synaptic connections to the same neuron are implemented by axonal connections that branch only locally, so that the additional wire length caused by the branching can be neglected.

A similar upper bound can be computed for the wire length, when one assumes $S = 1$:

$$WL(i) = \sum_{k=1}^{\infty} \sum_{J} \alpha k f(k) M_J p_{I \to J} e^{-\left(\frac{\alpha k}{\sigma}\right)^2} \tag{20}$$

$$= \alpha \sum_{J} M_J p_{I \to J} \sum_{k=1}^{\infty} k f(k) \gamma^{k^2} \tag{21}$$

, where

$$\gamma = e^{\left(\frac{\alpha}{\sigma}\right)^2}. \tag{22}$$

Replacing the function $f(k)$ yields:

$$WL(i) = 8\alpha \sum_{J} M_J p_{I \to J} \left( \sum_{k=1}^{\infty} k^2 \gamma^{k^2} \right). \tag{23}$$

We get an upper bound by using a standard estimate for the geometric series:

$$8\alpha \sum_{J} M_J p_{I \to J} \left( \sum_{k=1}^{\infty} k^2 \gamma^{k^2} \right) < 8\alpha \sum_{J} M_J p_{I \to J} \left( \sum_{k=1}^{\infty} k^2 \gamma^{k} \right) \tag{24}$$

$$= 8\alpha \sum_{J} M_J p_{I \to J} \left( \frac{\gamma(\gamma + 1)}{(1 - \gamma)^3} \right). \tag{25}$$

Note that this upper bound does not contain anymore an infinite sum.

For a more general upper bound where $S > 1$ one can consider $S$ RSNNs superimposed, resulting in:

$$WL(i) \leq 8\alpha S \sum_{J} M_J p_{I \to J} \left( \frac{\gamma(\gamma + 1)}{(1 - \gamma)^3} \right). \tag{26}$$

**Sparsity and wire length of RSNN samples:** Table 1 shows the average values of connection sparsity and the total wire length from RSNNs generated by the probabilistic skeletons for each tasks. The sparsity refers to the fraction of synaptic connections between neurons i and j that are realized, i.e. $m_{ij} > 0$..

| Task | # of neurons | Sparsity | Avg. # synapses | Wire length |
|---|---|---|---|---|
| Delayed pattern matching | 300 | 23.3% | 17.500 | 1.53m |
| Spike interval classification | 304 | 24.6% | 47,700 | 1.27m |
| Spike pattern classification | 148 | 37.8% | 30,500 | 0.595m |
| Contrast enhancement | 200 | 16.03% | 4,810 | 0.505m |
| Coincidence detection | 375 | 15.44% | 54,000 | 1.47m |
| Ant | 458 | 11.24% | 52,700 | 0.79m |

**Table 1:** Number of neurons, average connection sparsity, average number of synapses and wire length of RSNNs emerging from the probabilistic skeletons on the different tasks. On the contrast enhancement and the coincidence detection task the RSNNs consisting of 25 minicolumns were considered. Note that the number of synapses in an RSNN will vary between different samples from the same probabilistic skeleton. Hence, the average over 5 RSNN samples has been considered for this value.

#### 1.9 Population coding

Each continuous-valued input value is encoded by the spiking activity of an excitatory population of $N_{\text{pop}}$ input neurons. An input variable $x$ can only take values in the bounded interval $(a, b] \subset \mathbb{R}$. We define points $a = x_1, \ldots, x_M = b$ such that the subintervals $(x_k, x_{k+1}],\ k = 0, \ldots, N_{\text{pop}}$ are of equal lengths $\frac{b-a}{N_{\text{pop}}-1}$ and disjointly overlap the interval. The neurons are chosen such that there is a positive linear dependency between the input values and the spatial position of the neurons. On these subintervals gaussian density functions are used to model the probability that a neuron $i$ spikes for a given input $x \in (a, b]$ by first defining

$$h_k(x) = \frac{1}{\sigma_{pop}\sqrt{2\pi}} \exp\left\{-\frac{1}{2}\left(\frac{x - x_k}{\sigma_{pop}}\right)^2\right\} \tag{27}$$

for $k = 1, \ldots, N_{\text{pop}}$, where $\sigma_{pop} \in \mathbb{R}$. In the experiments it is chosen to be $\sigma_{pop} = \frac{x_{k+1}-x_{k-1}}{2}$, such that $\sim 68.27\%$ of spiking activity for this type happens in the interval $(x_{k-1}, x_{k+1}]$ and $\sim 95.45\%$ happens in the interval $(x_{k-2}, x_{k+2}]$ for suitable $k$. Since this is modeled by a density function it is necessary to normalize the values of $h_k$ to obtain spike probabilities for each neuron. A schematic plot of the population coding is given in Figure S5.

#### Illustration of population coding.

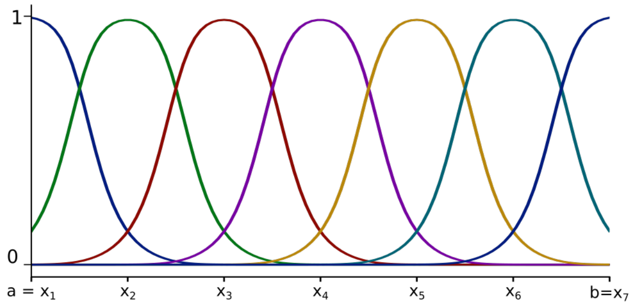

**Figure S5:** Illustration of population coding.

The spike train $z_i : \mathbb{R} \to \mathbb{R}$ for an input neuron $i$ and an input value $x$ is therefore given as

$$z_i(t) = \begin{cases} 1 & \text{with probability } \frac{h_i(x)}{\max\limits_{x \in [a,b]} h_i(x)} \\ 0 & \text{else.} \end{cases} \tag{28}$$

#### 1.10 Hardware specifications

Most of our experiments were conducted using 2 × AMD EPYC Rome 7402 CPUs with 2×24 cores and 2.8 GHz, 4 × NVIDIA A100 GPU and 4 × 40 GB HBM2e (Juwels Booster)

207 # 2 Supplementary figures

208 ## Additional spike rasters for the computations on spike times

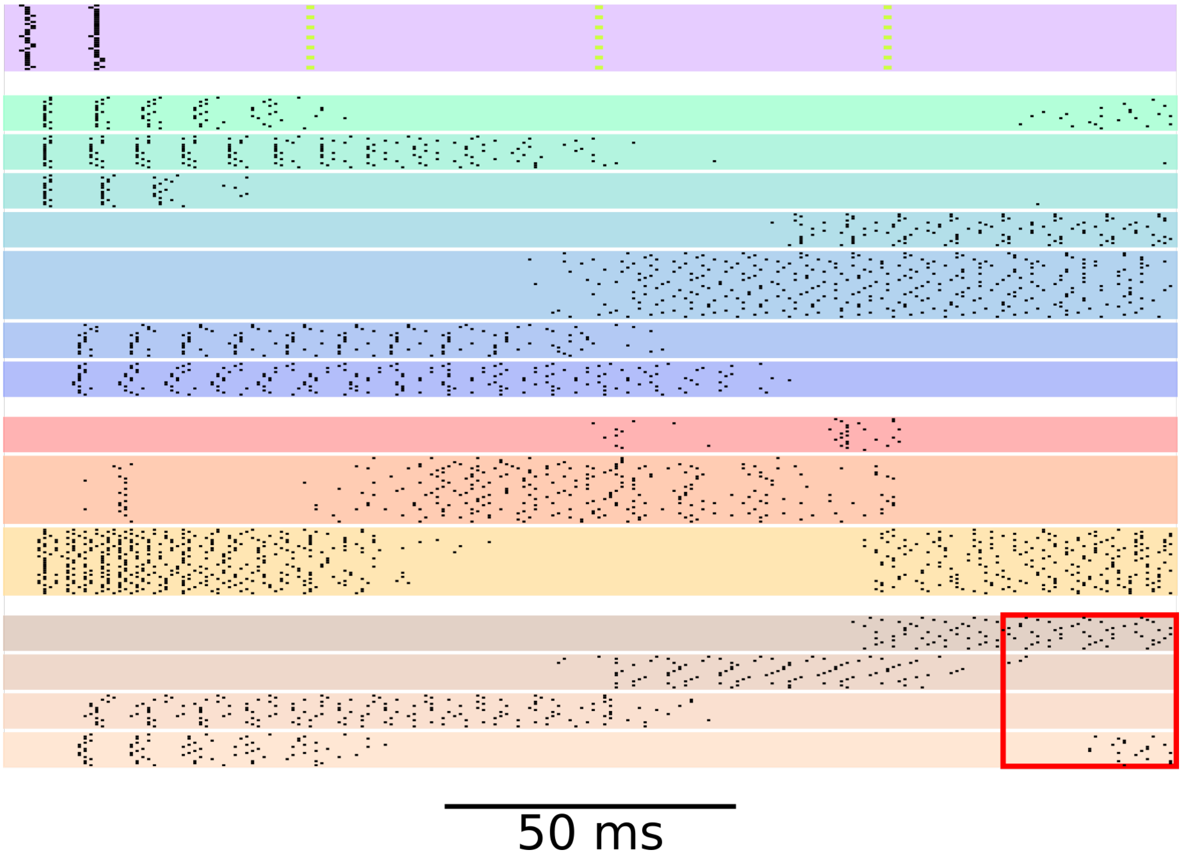

**Figure S6:** Firing activity of the same RSNN as in Fig. 3 d, for the case where the delay between the two input spike waves was between 0 and 50 ms. The RSNN correctly assigned the input to class 1.

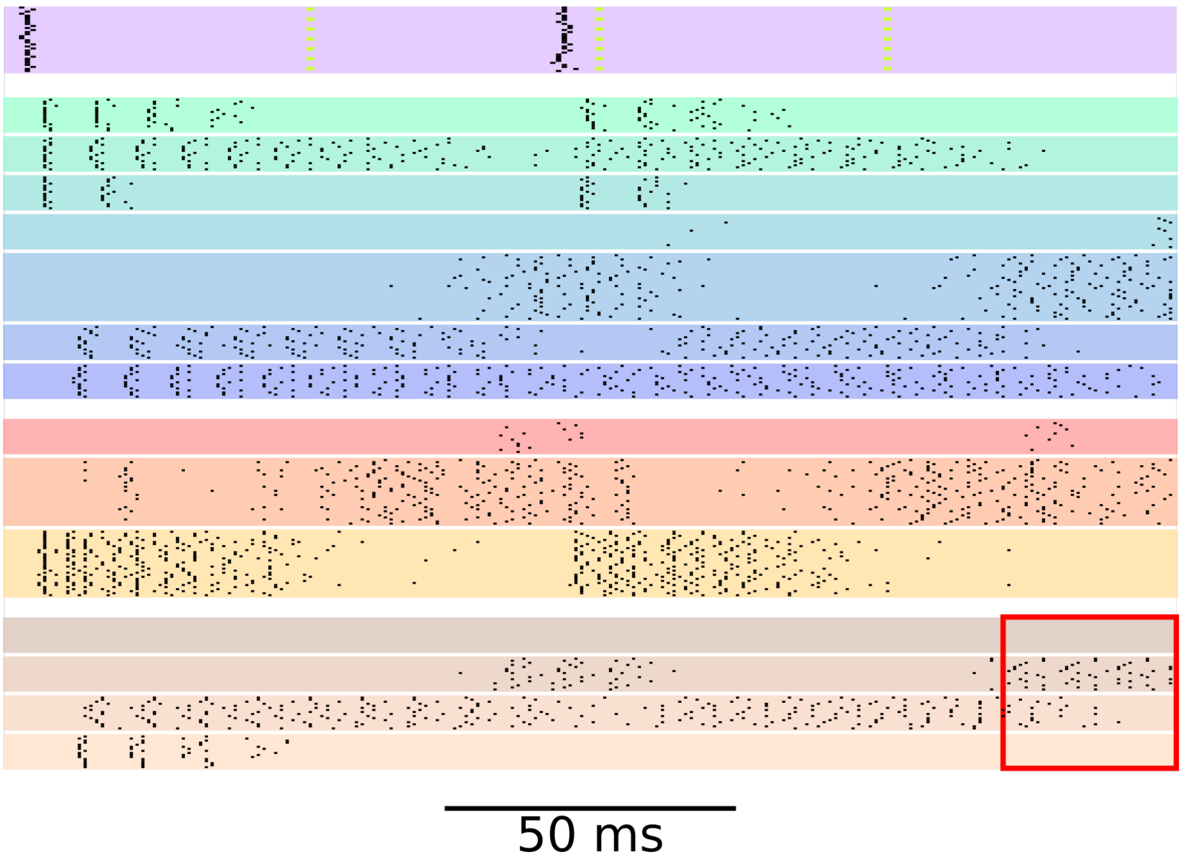

**Figure S7:** Firing activity of the same RSNN as in Fig. 3 d, for the case where the delay between the two input spike waves was between 50 and 100 ms. The RSNN correctly assigned the input to class 2.

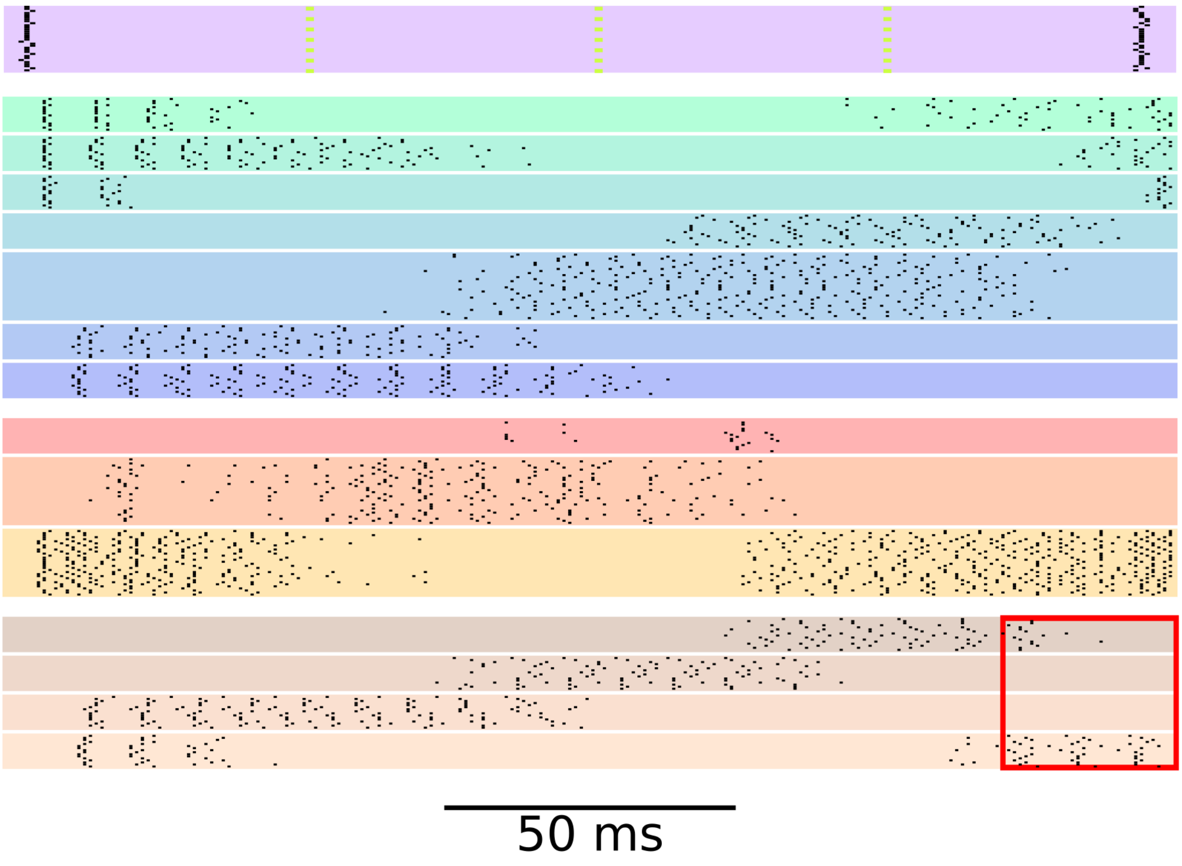

**Figure S8:** Firing activity of the same RSNN as in Fig. 3 d, for the case where the delay between the two input spike waves was between 150 and 200 ms. The RSNN correctly assigned the input to class 4.

#### Connection probabilities for ant, including connections of input and output neuron types

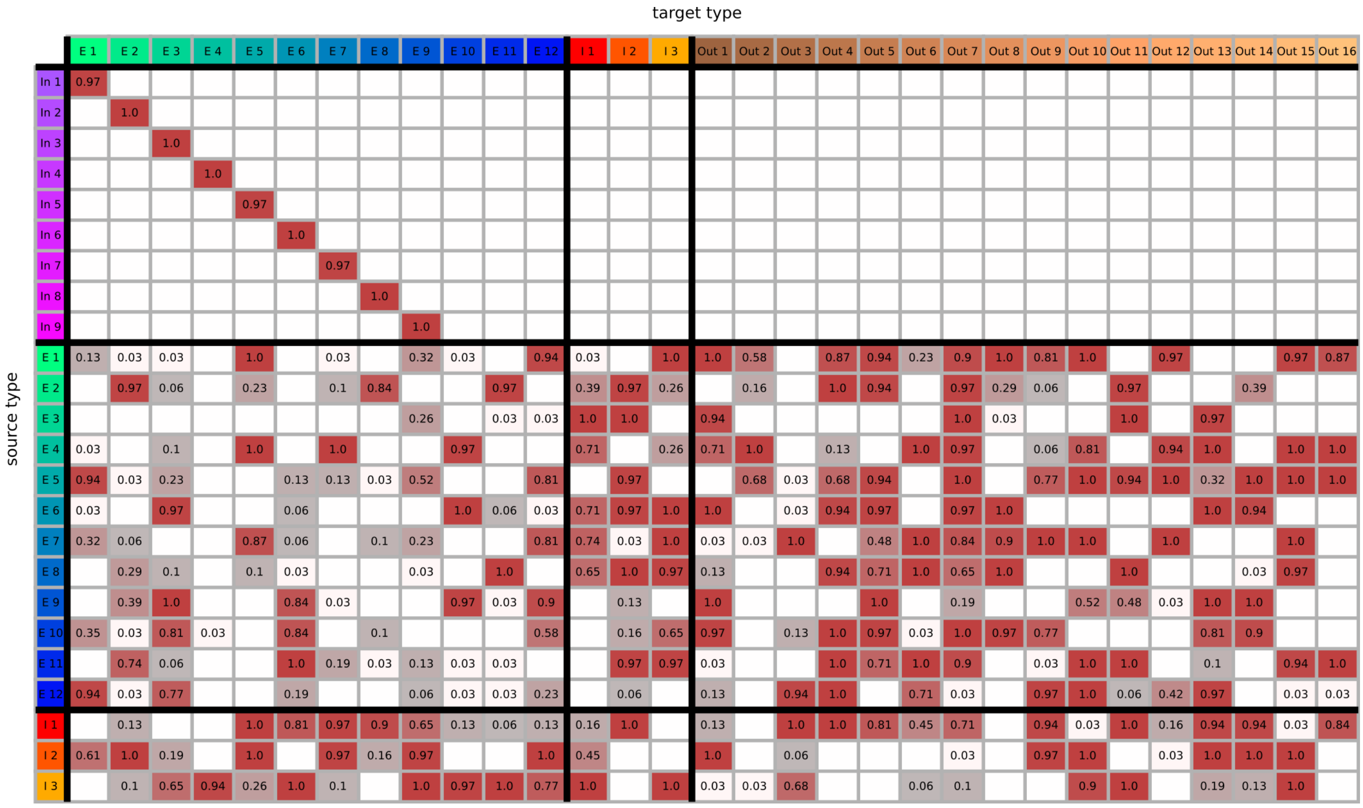

**Figure S9:** Full connection probabilities of the probabilistic skeleton on the ant task. Note, that input types were restricted to only connect to one exclusive recurrent type.
